# An N-terminal domain specifies developmental control by the SMAX1-LIKE family of transcriptional co-repressors in *Arabidopsis thaliana*

**DOI:** 10.1101/2024.05.12.593779

**Authors:** Sun Hyun Chang, Wesley J. George, David C. Nelson

**Author notes:** **Corresponding author:** David C. Nelson. **The author(s) responsible for distribution of materials integral to the findings presented in this article is:** David C. Nelson.

## Abstract

SMAX1-LIKE (SMXL) proteins are transcriptional co-repressors that regulate many aspects of plant growth and development. Proteins from the SMAX1- and SMXL78-clades of this family are targeted for degradation after karrikin or strigolactone perception, triggering downstream responses. We investigated how SMXL proteins control development. *SMXL7* can partially replicate *SMAX1* function in seeds and seedlings, but *SMAX1* cannot replace *SMXL7* in shoot branching control. Therefore, the distinct roles of these genes arise from differences in protein activity more than expression. Analysis of chimeras and domain deletions of SMAX1 and SMXL7 proteins revealed that an N-terminal domain is necessary and sufficient to specify developmental functions. We screened 158 transcription factors for interactions with SMAX1. The N-terminal domain is necessary and/or sufficient for the majority of candidate interactions. These discoveries enable cross-wiring of karrikin and strigolactone control of plant development and lay a foundation for understanding how SMXL proteins evolved functional differences.

## INTRODUCTION

Plant hormones control growth, development, and responses to the environment through regulation of transcriptional networks (Yin et al. 2023). Several plant hormones, including auxin, jasmonate, gibberellin, and strigolactone, initiate downstream responses through hormone-triggered polyubiquitination and degradation of transcriptional regulatory proteins (Blázquez et al. 2020). Strigolactones (SLs), for example, promote protein-protein interactions between an ɑ/ꞵ-hydrolase receptor, DWARF14 (D14)/DECREASED APICAL DOMINANCE2 (DAD2), an F-box protein within an SCF (Skp1-Cullin-F-box) E3 ubiquitin ligase complex, MORE AXILLARY GROWTH2 (MAX2), and a subset of proteins within the SUPPRESSOR OF MAX2 1 (SMAX1)-LIKE (SMXL)/DWARF(D53) family. The associated SMXL proteins are then polyubiquitinated by SCF^MAX2^ and rapidly destroyed by the 26S proteasome (Jiang et al. 2013; Wang et al. 2015; Zhou et al. 2013; Soundappan et al. 2015; de Saint Germain et al. 2016; Yao et al. 2016; Hamiaux et al. 2012). SMXL proteins are thought to function as transcriptional co-repressors, as they interact with TOPLESS (TPL)/TPL-RELATED (TPR) proteins via one or more EAR (Ethylene-responsive element binding factor-associated Amphiphilic Repression) motifs (Soundappan et al. 2015; Ma et al. 2017; Jiang et al. 2013; Wang et al. 2015; Liang et al. 2016). Thus, SMXL degradation initiates downstream responses through the release of transcriptional repression.

Proteins within the SMXL family have diversified to regulate different developmental processes and to be regulated, in turn, by different signaling mechanisms. In angiosperms, SMXL proteins are grouped into four phylogenetic clades: aSMAX1, SMXL39, aSMXL4, and SMXL78 (Walker et al. 2019). aSMAX1-clade proteins, represented by SMAX1 and SMXL2 in *Arabidopsis thaliana* or OsSMAX1 in *Oryza sativa* (rice), regulate seed germination, seedling photomorphogenesis (or in rice, mesocotyl elongation in the dark), root hair density and elongation, drought tolerance, and symbiotic interactions with arbuscular mycorrhizal fungi (Stanga et al. 2013; Feng et al. 2022; Villaécija-Aguilar et al. 2022; Park et al. 2022; Bursch et al. 2021; Carbonnel et al. 2020a; Bunsick et al. 2020; Choi et al. 2020; Zheng et al. 2020). SMAX1 and SMXL2 are targeted for degradation in an SCF^MAX2^-dependent manner by a paralog of D14, KARRIKIN INSENSITIVE2 (KAI2)/HYPOSENSITIVE TO LIGHT (HTL) (Khosla et al. 2020a; Wang et al. 2020b; Zheng et al. 2020). KAI2 putatively mediates responses to a metabolite of karrikins (KARs), butenolide molecules found in smoke, as well as an undiscovered endogenous compound(s) known as KAI2 ligand (KL) (Waters and Nelson 2022). Diversification of KAI2 proteins in some lineages has led to selective recognition of different KARs or alternative ligands such as SLs and (–)-germacrene D (Conn et al. 2015; Toh et al. 2015; Tsuchiya et al. 2015; Stirling et al. 2024; Martinez et al. 2022; Guercio et al. 2022; de Saint Germain et al. 2021; Carbonnel et al. 2020b; Sun et al. 2020; Conn and Nelson 2015). SMAX1 and SMXL2 can also be targeted by D14– SCF^MAX2^ when SLs are sufficiently abundant (Li et al. 2022; Wang et al. 2020b). SMXL78-clade proteins, represented by SMXL6, SMXL7, and SMXL8 in *Arabidopsis* and D53 in rice, control axillary branching or tillering, secondary growth, leaf elongation, internode elongation, and more (Jiang et al. 2013; Zhou et al. 2013; Agusti et al. 2011; Liang et al. 2016; Wang et al. 2020a; Yang et al. 2020; de Saint Germain et al. 2013; Li et al. 2020). These proteins are specifically targeted by D14 and not by KAI2 (Wang et al. 2015; Khosla et al. 2020a; White et al. 2022). Finally, SMXL3- and aSMXL4-clade proteins, represented by SMXL3, SMXL4, and SMXL5 in *Arabidopsis*, regulate phloem development and anthocyanin abundance (Cho et al. 2018; Wallner et al. 2017, 2023; Wu et al. 2017; Li et al. 2024). Unlike other members of the SMXL family, these proteins are not targeted for degradation by SCF^MAX2^, putatively due to loss of a P-loop or Arg-Gly-Lys-Thr (RGKT) motif (Wallner et al. 2017). In addition to imposing transcriptional regulation on its own, SMXL5, and perhaps its similarly stable homologs, attenuates SL signaling by inhibiting degradation of SMXL7 (Li et al. 2024).

The diverse functions of SMXL proteins in plants raise the largely unanswered questions of what genes do SMXL proteins regulate and how do they do so? SMXL proteins are distantly related to a ClpB-type heat shock protein, HSP101, that forms hexameric ATPase complexes involved in solubilizing protein aggregates (Gallie et al. 2002; Stanga et al. 2013). SMXL proteins are composed of a Clp N-terminal domain, a degenerate ATPase domain (D1), a middle region (M), and a C-terminal, degenerate ATPase domain (D2) (Zhou et al. 2013; Wang et al. 2011; Khosla et al. 2020a). Structure-function analyses have revealed roles for several SMXL protein features. The D1M region confers specificity for SMXL interactions with D14 or KAI2 (Khosla et al. 2020a). The D2 domain contains the aforementioned RGKT and EAR motifs (Jiang et al. 2013; Soundappan et al. 2015; Wang et al. 2015; Liang et al. 2016; Ma et al. 2017). D2 is necessary for degradation of SMXL proteins, but is not sufficient except when full-length SMXL proteins are also present (Khosla et al. 2020a). This is likely due to multimeric SMXL complexes that are formed at least in part through interactions at the C-terminus (Khosla et al. 2020a). The EAR motif also likely contributes to stabilization of multimeric SMXL complexes (Ma et al. 2017; Li et al. 2024).

Only a few studies have identified direct genomic targets of SMXL proteins and partner proteins that putatively guide SMXL associations with DNA (Wang et al. 2020a; Song et al. 2017; Hu et al. 2020; Xu et al. 2023; Kim et al. 2022; Fang et al. 2020). ChIP-seq analysis of SMXL6 revealed 729 candidate target sites in the *Arabidopsis* genome, although there was little overlap with 401 SL-responsive genes (Wang et al. 2020a). Genes directly targeted by SMXL6 include *BRC1*, *SMXL2*, *SMXL6*, *SMXL7*, and *SMXL8 (Wang et al. 2020a)*. Unexpectedly, SMXL6 and SMAX1 can bind DNA directly; both proteins recognize the motif 5’-ATAACAA-3’ or 5’-TTGTTAT-3’ (Wang et al. 2020a; Xu et al. 2023). In rice, D53 interacts with the transcription factors BRI1-EMS SUPPRESSOR1 (OsBES1), GROWTH REGULATORY FACTOR4 (OsGRF4), REDUCED LEAF ANGLE1 (OsRLA1), and IDEAL PLANT ARCHITECTURE1 (OsIPA1) (Song et al. 2017; Hu et al. 2020; Sun et al. 2023; Fang et al. 2020). In *Arabidopsis*, SMAX1 interacts with phytochrome B and DELLA proteins (Park et al. 2022; Kim et al. 2022; Xu et al. 2023). Therefore, transcriptional regulation by SMXL proteins may arise from a combination of binding specific cis-regulatory motifs as well as associating with transcription factors (TFs). Here, we investigated the molecular basis of transcriptional control by SMXL proteins.

## RESULTS

### SMAX1 and SMXL7 are not interchangeable

Differential expression of genes in the *SMAX1*- and *SMXL78*-clades occurs in many tissue types and developmental stages of *Arabidopsis thaliana*, although in some cases both types of genes show similar expression (Stanga et al. 2013; Soundappan et al. 2015). For example, *SMAX1*-clade transcripts are enriched in seeds and emerging seedlings, while *SMXL78*-clade transcripts are enriched in the roots and apices of older seedlings; however, both types of transcripts are abundant in leaves and floral tissues (Supplemental Fig. S1) (Klepikova et al. 2016).

This led us to investigate whether the different roles of the *SMAX1-* and *SMXL78-*clades in *Arabidopsis* development are due to their expression patterns. We focused on *SMAX1* and *SMXL7* as representative members of each clade because they generally showed the highest expression (Supplemental Fig. S1). We performed a promoter-swapping experiment in which we tested whether *SMXL7* expressed under the control of a *SMAX1* promoter (*SMAX1pro::SMXL7*) could rescue the *smax1-2 smxl2-1* (hereafter, *smax1,2*) mutant. *SMAX1pro::SMAX1* fully rescued the hypocotyl elongation of *smax1,2* seedlings grown under dim red light. In contrast, *SMAX1pro::SMXL7* partially rescued hypocotyl growth and inhibited cotyledon expansion (Fig. 1 A and B). Similarly, while *SMAX1pro::SMAX1* restored seed dormancy to *smax1,2*, *SMAX1pro::SMXL7* had a significantly weaker effect (Fig. 1C). This implies that *SMXL7* shares some function with *SMAX1* but the genes are not equivalent.

**Figure 1.**
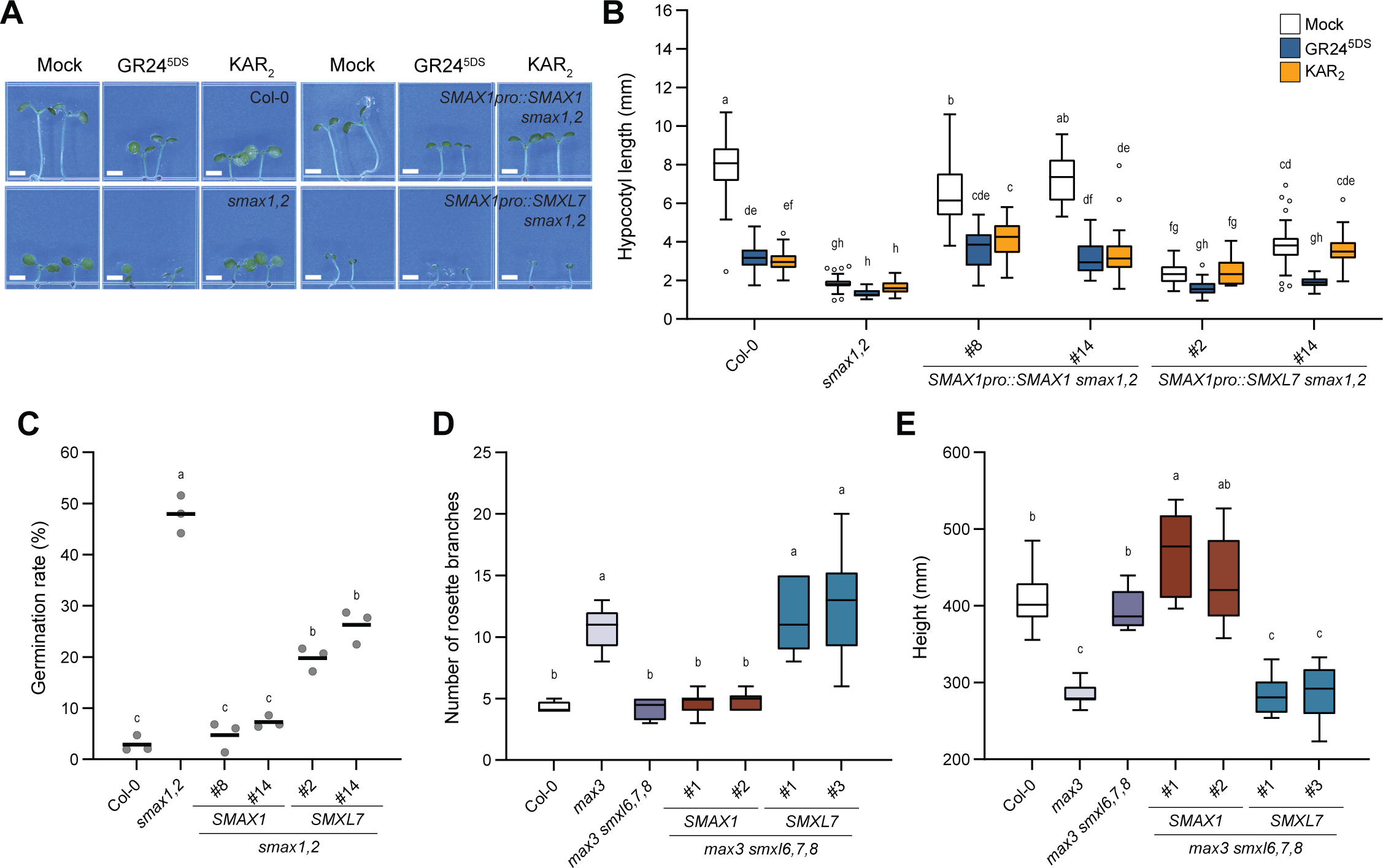
Differences in *SMAX1* and *SMXL7* functions are not due to expression alone. (*A*) Hypocotyl length of Col-0, *smax1,2*, and transgenic seedlings grown 6 d in red light expressing *SMAX1pro::SMAX1* or *SMAX1pro::SMXL7* in the *smax1,2* background. Seedlings were treated with mock, 1 μM KAR_2_, or 1 μM GR24^5DS^ (n>20). (*B*) Representative image of the seedlings used in *A*. Scale bar = 2 mm. (*C*) Germination of transgenic lines expressing *SMAX1* and *SMXL7* under the *SMAX1* promoter in *smax1,2*. (n=3, >50 seeds per replicate) (*D*) Rosette branch numbers of 8-week-old Col-0, *max3*, *max3 smxl6,7,8*, and *SMXL7pro::SMAX1* or *SMXL7pro::SMXL7* in *max3 smxl6,7,8* (n>9). (*E*) Height of plants in *D*. Boxplots indicate mean with quartiles and Tukey’s whiskers; open symbols are outlier points that fall beyond the range of the whiskers. Letters indicate groups with significant differences (*P*<0.05, two-way ANOVA in *B*, or one-way ANOVA in *C-E*, followed by Tukey’s multiple comparisons test).

Regulation of SMAX1 and SMXL7 in these transgenic seedlings was consistent with prior studies (Fig. 1B). Hypocotyl elongation of *SMAX1pro::SMAX1 smax1,2* seedlings was inhibited by KAR_2_ and a synthetic SL, GR24^5DS^, implying KAI2- and D14-induced degradation of SMAX1, respectively. *SMAX1pro::SMXL7 smax1,2* seedlings, however, were responsive to GR24^5DS^ only, consistent with D14-specific targeting of SMXL7 for degradation (Fig. 1B). Therefore, even though the *SMXL78* clade has little or no control of hypocotyl elongation in *Arabidopsis* (Soundappan et al. 2015; Li et al. 2022), regulation of misexpressed *SMXL7* is intact in seedlings.

We then tested the converse situation: could *SMAX1* replace a *SMXL78*-clade deficiency when expressed under the control of a *SMXL7* promoter? To avoid D14-induced degradation of SMAX1 (Li et al. 2022; Wang et al. 2020b), which might reduce the effectiveness of a *SMXL7pro::SMAX1* transgene, and to maximize the phenotypic differences between rescued and non-rescued lines, we introduced *SMXL7pro::SMAX1* into *max3 smxl6,7,8*. This quadruple mutant is SL-deficient, but also has constitutive SL responses. *SMXL7pro::SMXL7* rescued the axillary branching and shoot height phenotypes of *max3 smxl6,7,8* to those seen in the *max3* single mutant. In contrast, *SMXL7pro::SMAX1* did not affect either shoot phenotype (Fig. 1 D and E). Altogether these observations demonstrated that the unique functions of *SMAX1* and *SMXL7* are not simply a consequence of their expression patterns. Therefore, SMAX1 and SMXL7 proteins likely regulate distinct developmental processes by regulating different sets of genes, for example through selective interactions with transcription factor partners.

### An N-terminal “output” domain specifies developmental control by SMAX1 and SMXL7

To determine which part of SMXL proteins specifies their roles in development, we performed a structure-function analysis. Our first strategy was to swap major domains of SMAX1 and SMXL7 proteins and test the functions of the resulting chimeras (hereafter, SMXLχ, where the Greek letter χ represents chimera). Domains in SMAX1 and SMXL7 have three conserved globular regions connected by less conserved, often intrinsically disordered regions (IDRs) of variable lengths (Supplemental Fig. S2 and S7A) (Khosla et al. 2020a; Tal et al. 2022; Temmerman et al. 2022). Therefore, to keep the swapped domains in a near-native context within the broader protein structure, we adjusted the previously defined domain boundaries of SMAX1 and SMXL7 to end at nearby, highly conserved residues (Supplemental Fig. S2). We also considered a prediction of SMAX1 protein structure created by AlphaFold2 (Jumper et al. 2021). This model suggested that our initial C-terminal boundary for the SMAX1_N_ domain at aa 158 is located in the middle of the ninth alpha helix (Supplemental Fig. S1). This led us to also test a longer, 210-aa version of the SMAX1_N_ domain that encompasses the predicted globular N-terminal region and part of a putative IDR that follows it. We created a series of reciprocal swaps of the N, D1 and M (D1M), and D2 domains from SMAX1 and SMXL7 (Fig. 2A). (The three numbers following SMXLχ in each chimera name indicate the source of the N, D1M, and D2 domains, respectively.) To validate the expression and correct subcellular localization of the SMXLχ proteins, we created N-terminal fusions with eYFP and transiently expressed each construct in *Nicotiana benthamiana* leaves. All eYFP-SMXLχ proteins produced nuclear-localized fluorescence that was consistent with the localization of wild-type SMAX1 and SMXL7 (Supplemental Fig. S3).

**Figure 2.**
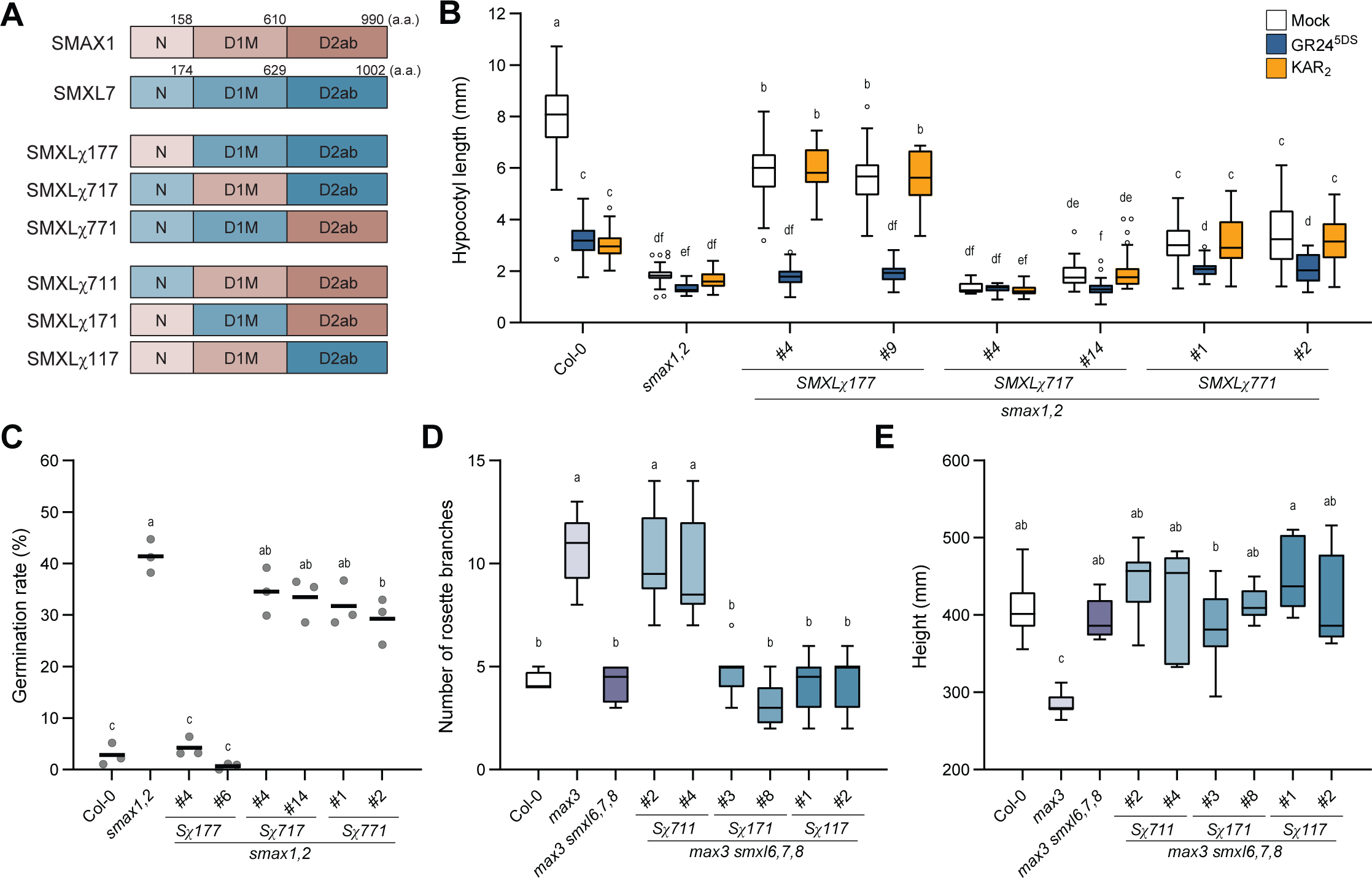
Chimera analysis implicates N domain of SMXL proteins in developmental control. (*A*) Schematic of N, D1M, and D2 domain boundaries of SMAX1, SMXL7, and chimeric SMXL proteins. (*B*) Hypocotyl length of red light-grown seedlings treated with mock, 1 μM KAR_2_, or 1 μM GR24^5DS^ (n>20). (*C*) Germination Col-0, *smax1,2*, and transgenic lines expressing *SMXL!177*, *SMXL!717*, and *SMXL!771* under the *SMAX1* promoter in *smax1,2.* (n=3, >50 seeds per replicate) (*D*) Rosette axillary branch numbers in Col-0, *max3*, *max3 smxl6,7,8*, and transgenic lines expressing *SMXL!711*, *SMXL!171*, and *SMXL!117* under control of the *SMXL7* promoter in *max3 smxl6,7,8* (n>9). (*E*) Height of the plants in *D*. Boxplots indicate mean with quartiles and Tukey’s whiskers; open symbols are outlier points that fall beyond the range of the whiskers. Letters indicate groups with significant differences (*P*<0.05, two-way ANOVA in *B*, or one-way ANOVA in *C-E*, followed by Tukey’s multiple comparisons test). S*!*, abbreviation for SMXL*!*.

We then tested whether the SMXLχ proteins could rescue *smax1,2* or *max3 smxl6,7,8* when expressed under the control of *SMAX1* or *SMXL7* promoters, respectively. *SMAX1pro::SMXLχ177* mostly rescued the short hypocotyl phenotype of *smax1,2* (Fig. 2B). In contrast, *SMAX1pro::SMXLχ771* rescued hypocotyl elongation weakly, similar to *SMAX1pro::SMXL7*, and *SMAX1pro::SMXLχ717* had no effect (Fig. 1A and 2B). This suggested that SMAX1_N_ is sufficient to specify control of hypocotyl growth. Furthermore, because *SMXLχ711* had limited ability to rescue *smax1,2* hypocotyl elongation, the N domain may be necessary for SMAX1 function (Supplemental Fig. S4B). In case SMXL7 turnover limits its effectiveness in *smax1,2*, we tested an Arg-Gly-Lys-Thr (RGKT) deletion mutant that is resistant to SL-induced degradation (Zhou et al. 2013; Jiang et al. 2013; Wang et al. 2015; Soundappan et al. 2015). Like wild-type SMXL7, SMXL7^ΔRGKT^ recovered hypocotyl elongation of *smax1,2* only partially (Supplemental Fig. S4 C and D). *SMXLχ177*^ΔRGKT^ was more effective at rescuing hypocotyl length, and *SMXLχ1_210_77*^ΔRGKT^ even moreso (Supplemental Fig. S4 C and D).

Hypocotyl elongation of *SMAX1pro::SMXLχ177* and *SMAX1pro::SMXLχ771 smax1,2* seedlings was inhibited by GR24^5DS^ treatment, but not by KAR_2_ (Fig. 2B). This suggests that these chimeric proteins, which share the D1M domain from SMXL7, are targeted for degradation by D14 but not KAI2. It provides further evidence that the D1M domain specifies SMXL interactions with D14 or KAI2 receptors (Khosla et al. 2020a) and also demonstrates cross-wiring of the KAR and SL signaling systems. Unexpectedly, we also saw hypocotyl elongation responses to GR24^5DS^ in *SMXL7^ΔRGKT^*, *SMXLχ177^ΔRGKT^*, and *SMXLχ1_210_77^ΔRGKT^ smax1,2* lines, which may indicate that the RGKT deletion does not confer degradation resistance in this background (Supplemental Fig. S4 C and D). In the *max3 smxl6,7,8* background, however, SMXL7^ΔRGKT^ functioned as expected for a hypermorphic protein; *SMXL7pro::SMXL7^ΔRGKT^* had a stronger effect than *SMXL7pro::SMXL7*, producing shoot branching and height phenotypes that were even more dramatic than *max3* (Supplemental Fig. S5).

We observed similar results in germination assays of the *SMXLχ smax1,2* transgenic lines. *SMAX1pro::SMXLχ177* restored seed dormancy to *smax1,2,* but *SMAX1pro::SMXLχ717* and *SMAX1pro::SMXLχ771* did not affect germination significantly (Fig. 2C). Likewise, *SMXLχ177*^ΔRGKT^ and *SMXLχ1_210_77*^ΔRGKT^ rescued seed dormancy to a greater degree than *SMXL7*^ΔRGKT^ (Supplemental Fig. S4D).

We next investigated whether the N domain of SMXL7 specifies control of axillary branching and shoot height (Fig. 2 D and E). *SMXL7pro::SMXLχ711* restored the axillary branching of *max3 smxl6,7,8* to the level of *max3*, as we had observed for wild-type *SMXL7* (Fig. 1D and 2D). In contrast, *SMXL7pro::SMXLχ171* and *SMXL7pro::SMXLχ117* did not affect axillary branching (Fig. 2D). This suggests that SMXL7_N_ is sufficient to specify control of axillary branching. *SMXL7pro::SMXLχ177* also did not affect the axillary branching phenotype of *max3 smxl6,7,8* (Supplemental Fig. S5A), implying that SMXL7_N_ is necessary for axillary branching control. Interestingly, none of the chimeras affected the height of *max3 smxl6,7,8* plants (Supplemental Fig. S5B). Therefore, regions of SMXL7 in addition to the N domain may be required to control shoot height. As well as having increased axillary branching from the rosette (i.e. primary branches), *max3* has excess secondary branching from cauline nodes (Booker et al. 2004). We found that *SMXL7pro::SMXLχ711* did not rescue secondary branching (Supplemental Fig. S5 C and D). Therefore, SMXL7_N_ is only sufficient for regulation of rosette axillary branching.

Collectively, these results indicate that the N domains of SMAX1 and SMXL7 play crucial roles in specifying downstream signaling outputs. However, other domains may also contribute to the distinct functions of these proteins, in particular for SMXL7.

### SMAX1_N_ is necessary and sufficient for regulating early development

To further investigate the role of the N domain in developmental regulation by SMAX1 and SMXL7, we performed a deletion analysis. We fused an N-terminal nuclear localization signal (NLS) from simian virus 40 (SV40) to all truncated proteins in order to maintain the correct subcellular localization (Fig. 3A). *SMAX1pro::SMAX1ΔN* failed to rescue *smax1,2* hypocotyl elongation or seed dormancy (Fig. 3 B and C). Similarly, *SMXL7pro::SMXL7ΔN* did not rescue the axillary branching or shoot height phenotypes of *max3 smxl6,7,8* (Fig. 3D and E). Therefore, the N domain is necessary for SMAX1 and SMXL7 functions.

**Figure 3.**
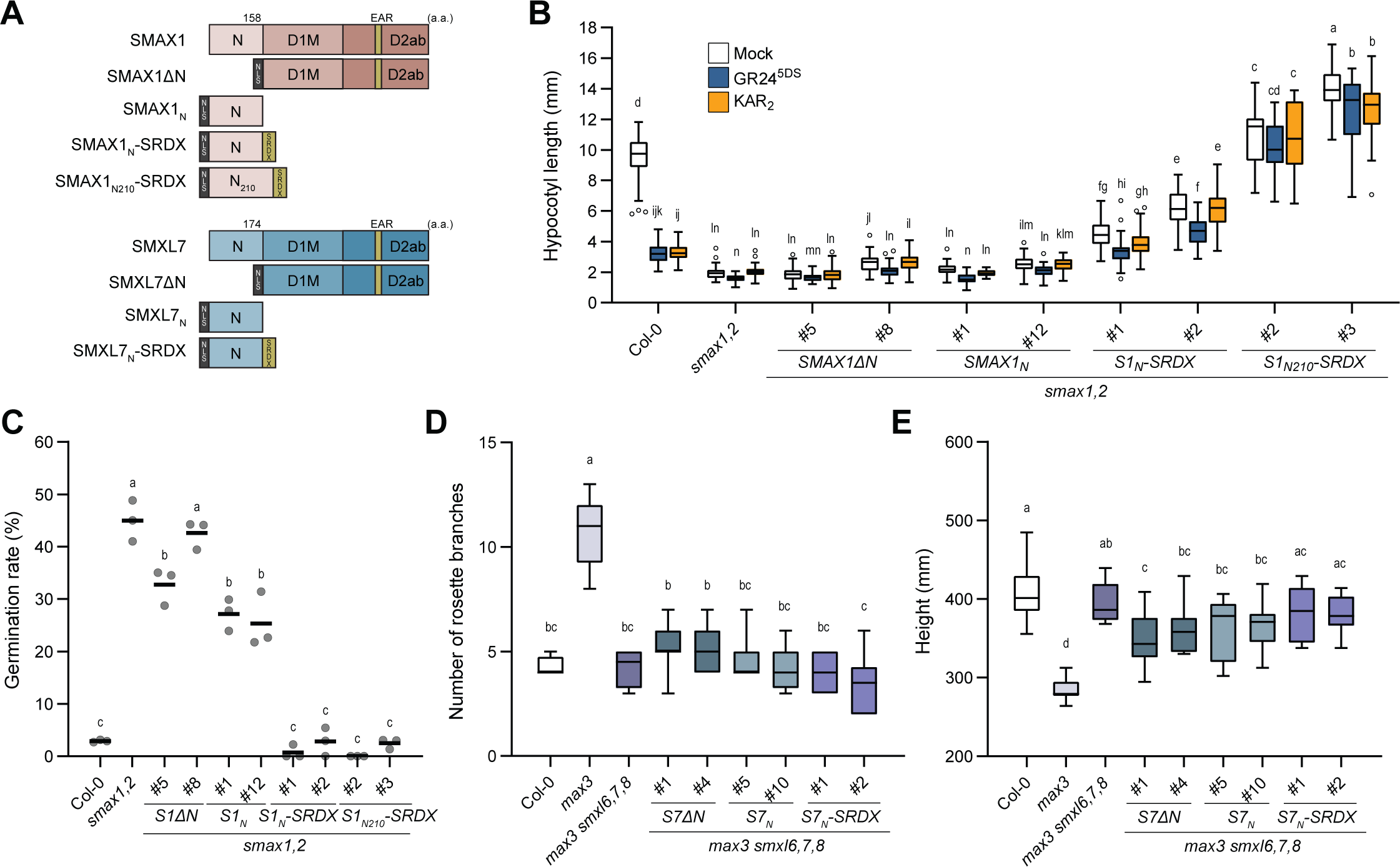
SMAX1_N_ specifies control of *Arabidopsis* germination and seedling growth. (*A*) Schematic representation of truncated SMAX1 and SMXL7 proteins. GFP-SRDX was used as a negative control. black, SV40 NLS added to the N-termini of truncated SMXL proteins. yellow, EAR or SRDX motifs. (*B-C*) Hypocotyl length of red light-grown seedlings treated with mock, 1 μM KAR_2_, or 1 μM GR24^5DS^ (*B*, n>20) and germination (*C*, n=3, >50 seeds per replicate) of Col-0, *smax1,2*, and transgenic lines expressing *SMAX1ΔN, SMAX1_N_, SMAX1_N_-SRDX,* and *SMAX1_N210_-SRDX* under control of the *SMAX1* promoter in *smax1,2*. (*D*) Rosette axillary branch numbers of Col-0, *max3*, *max3 smxl6,7,8*, and transgenic lines (n>9) expressing *SMXL7ΔN, SMXL7_N_,* and *SMXL7_N_-SRDX* (n>9). (*E*) height of plants in *D*. Boxplots indicate mean with quartiles and Tukey’s whiskers; open symbols are outlier points that fall beyond the range of the whiskers. Letters indicate groups with significant differences (*P*<0.05, two-way ANOVA in *B*, or one-way ANOVA in *C-E*, followed by Tukey’s multiple comparisons test). S1 or S7, abbreviations for SMAX1 or SMXL7.

We also tested SMAX1_N_ alone and found that it had no effect on *smax1,2* hypocotyl growth or germination (Fig. 3 B and C). This was not surprising, as SMXL functions are highly dependent on a C-terminal EAR motif(s) that facilitates interactions with TPL/TPR transcriptional co-repressors (Jiang et al. 2013; Wang et al. 2015; Liang et al. 2016; Ma et al. 2017; Soundappan et al. 2015; Li et al. 2024). To better mimic SMAX1 function, we fused SRDX, an artificial transcriptional repression domain based on EAR motif sequences (Hiratsu et al. 2003), to the C-terminus of SMAX1_N_. *SMAX1pro::SMAX1_N_- SRDX* moderately recovered hypocotyl elongation of *smax1,2* and restored seed dormancy (Fig. 3 B and C). A similar fusion with the longer version of SMAX1_N_ was more effective. *SMAX1_N210_-SRDX* robustly rescued *smax1,2*, causing hypocotyl elongation to exceed wild-type Col-0 (Fig. 3 B and C). This was not a consequence of SRDX alone, as *SMAX1pro::GFP-SRDX* had no effect on hypocotyl elongation of *smax1,2* and only affected seed germination weakly (Supplemental Fig. S6). These results demonstrate that SMAX1_N_, in particular the 210-aa version, is sufficient to specify regulation of germination and seedling growth. Notably, SMAX1_N210_-SRDX reconstitutes the function of full-length SMAX1 but not its regulation by SCF^MAX2^, as *SMAX1_N210_-SRDX smax1,2* seedlings were insensitive to KAR_2_ and GR24^5DS^ treatments (Fig. 3B).

We similarly tested the necessity and sufficiency of SMXL7_N_ for regulating SL responses. *SMXL7pro::SMXL7ΔN* did not rescue the axillary branching or shoot height phenotypes of *max3 smxl6,7,8*, supporting the necessity of the N domain for these functions. *SMXL7_N_* and *SMXL7_N_-SRDX* also failed to rescue *max3 smxl6,7,8* even though SMXL7_N_ was sufficient to confer axillary branching control to SMXLχ711. A longer N domain may be required to recapitulate SMXL7 function in an SRDX fusion or other domains may also be required to coordinate gene regulation.

### The N domain putatively mediates SMXL interactions with many transcription factors

SMXL6 and SMAX1 have been reported to bind to the same DNA motif (Wang et al. 2020a; Xu et al. 2023). This implies that additional factors are required to achieve distinct developmental outputs. We reasoned that SMAX1 may control different gene regulatory networks than SMXL7 through differential interactions with transcription factor (TF) protein partners. This led us to screen for potential TF partners of SMAX1 and SMXL7, simultaneously examining the importance of the N domain for such interactions.

We conducted yeast two-hybrid (Y2H) assays with 158 transcriptional regulators from an *Arabidopsis thaliana* TF library (Pruneda-Paz et al. 2014). The following criteria aided our selection of candidate TFs (Supplemental Table S1): 1) physical and/or genetic interactions with SMAX1 or SMXL78-clade proteins, 2) putative direct targets of SMXL6, 3) differential expression after GR24 treatment, and 4) association with seed development/germination, photomorphogenesis, root hair development, or leaf morphology (Wang et al. 2020a; Humphreys et al. 2023). We focused on identifying potential interactions between candidate TFs and SMAX1, which has been less characterized (Fig. 4, Supplemental Fig. S5). Potential SMAX1 interactors were then tested for interactions with SMXL7. The respective N domains (SMAX1_N_ and/or SMXL7_N_) or SMXL proteins lacking the N domain (SMAX1ΔN and/or SMXL7ΔN) were also tested to determine the basis of any positive Y2H interactions. If a TF interacted with a full-length SMXL protein but not the N domain alone, we tested whether it could interact with a longer version of the N domain (SMAX1_N210_ and/or SMXL7_N190_), as SMAX1_N210_-SRDX had proven more effective than SMAX1_N_-SRDX in transgenic plants.

**Figure 4.**
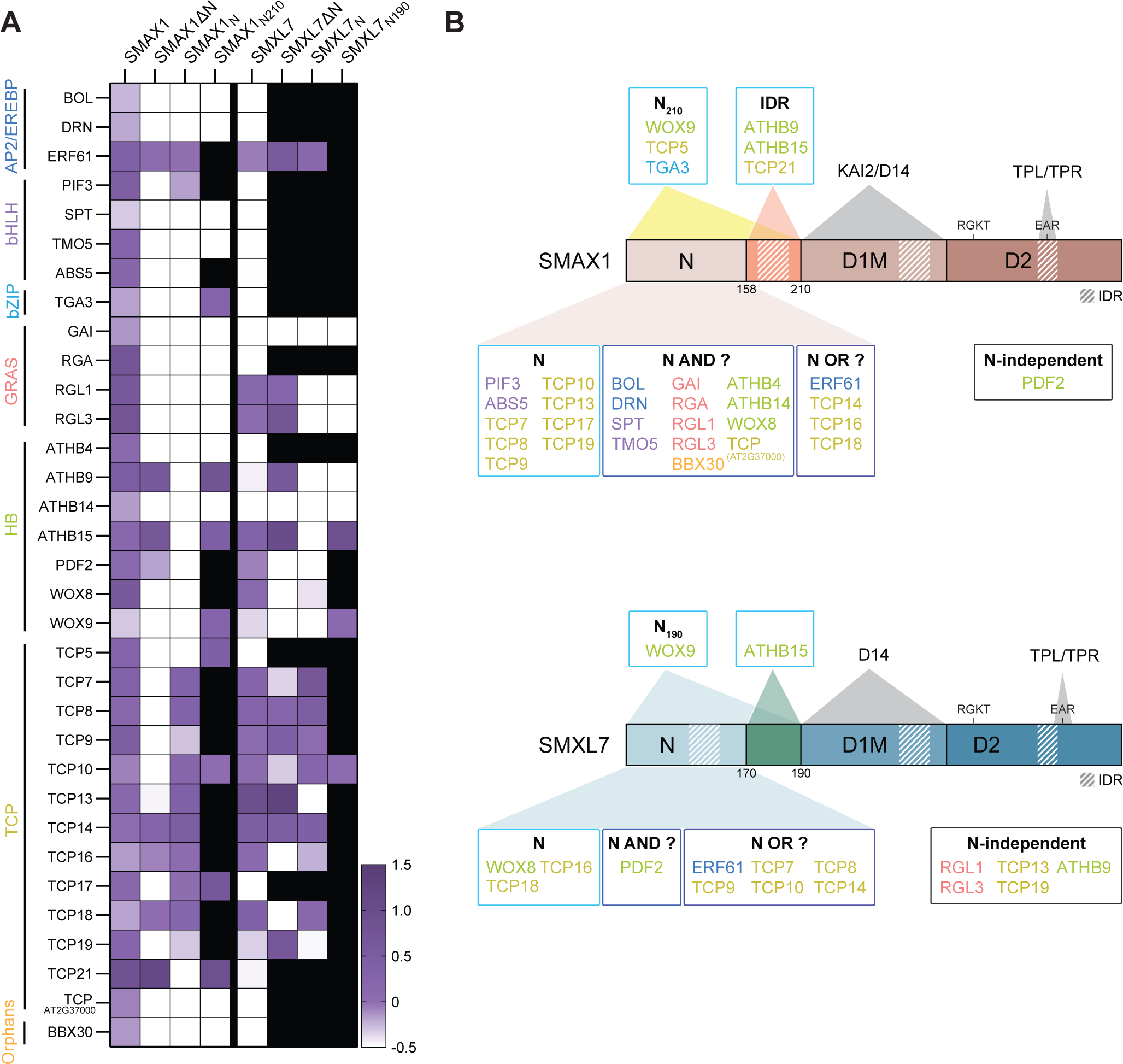
SMAX1_N_ is involved in most of the potential interactions with transcription factors. (*A*) Heatmap summarizing positive Y2H interactions between full-length and truncated SMAX1 or SMXL7 proteins with various TFs. Interaction level was quantified by comparing yeast growth on –LW and –LWH media and presented on a log_2_ scale. Interactions with values above -0.5 (purple) were considered positive, while below -0.5 (white) were considered non-interacting. Black boxes indicate untested combinations. (*B*) Schematic of the TFs that interact with the N-terminal domain of SMAX1 (top) or SMXL7 (bottom). TFs are grouped based on their interaction requirements with the SMAX1/SMXL7 N domain: “N”, TFs for which the N domain is both necessary and sufficient for interactions; “N AND ?”, TFs that require the N domain and an additional SMXL region(s) for interaction; “N OR ?”, TFs for which the N domain is sufficient but not necessary for interaction. hatched boxes, three longest predicted intrinsically disordered regions (IDRs) based on Supplemental Figure S7. TFs belonging to the same family are depicted in the same color in *A* and *B*.

We identified 33 TFs that showed positive Y2H interactions with full-length SMAX1. The majority of these interactions (25/33) required SMAX1_N_. The N domain was sufficient for interactions with eight of these proteins, and the longer 210-aa version of N was sufficient for interaction with three proteins. Fourteen proteins that required the N domain for interaction did not show interaction with any of the SMAX1 truncation proteins. This suggests that binding these TFs may require the N domain as well as another part of SMAX1. Of the remaining eight TFs that did not require the N domain for interaction with SMAX1, four could interact with the N domain alone and three putatively interacted with the 52-amino acid, IDR-containing region that distinguishes the long and short versions of SMAX1_N_. Altogether these observations support the importance of the N domain in transcriptional control by SMAX1 and identify a set of TFs that may be involved in developmental regulation by SMAX1. Notably, only 17 of the 33 SMAX1-interacting TFs also interacted with SMXL7 or its derivatives. Some of the differential Y2H interactions we observed with TFs may explain the unique roles of SMAX1 and SMXL7 in plant development.

## DISCUSSION

This study implicates the N domain of SMXL proteins as a major determinant of SMXL roles in plant growth and development. It remains to be seen whether the N domain is responsible for direct DNA-binding by SMXL proteins or only interactions with TFs. Because the N domain is more stable than full-length SMAX1, it may be more amenable to chromatin immunoprecipitation analysis and identification of SMAX1 protein partners via co-immunoprecipitation and tandem mass spectrometry. Further refinement of critical features within the N domain that specify developmental roles will also be useful in order to determine how SMXL proteins evolved different functions during the diversification of this family in the angiosperm lineage.

In future studies, it will be important to validate the candidate TF interactions with SMAX1 or SMXL7 through genetic and biochemical approaches. Of particular interest are TFs that may differentially interact with SMAX1 and SMXL7. Our screen suggested that SMAX1 may interact with multiple members of the TCP, HB, GRAS, bHLH, and AP2/EREBP families (Supplemental Table S1). TEOSINTE BRANCHED 1/CYCLOIDEA/PCF (TCP) family proteins are categorized into class I-PCF, class II-CIN, and class II-CYC/TB1 subclades. All class I TCPs that we tested, except for TCP21, interacted with both SMAX1 and SMXL7. Most of the tested class II TCPs also interacted with both SMAX1 and SMXL7, but TCP5 and TCP17 only interacted with SMAX1. TCP18/BRANCHED1, a key regulator of branching (Aguilar-Martínez et al. 2007; Wang et al. 2019), putatively interacted with SMAX1 and SMXL7 through the N domain. Regarding the GRAS family, we examined four of the five *Arabidopsis* DELLAs. GAI, RGL1, and RGL3 interacted with SMAX1 and SMXL7, but RGA showed SMAX1-specific interaction. The N domain was necessary but not sufficient for SMXL–DELLA interactions, consistent with a prior report that these DELLAs cannot interact with SMAX1 lacking its first 163 amino acids (Kim et al. 2022). In some cases, our Y2H results differed from expectations. For example, we did not observe clear SMAX1 or SMXL7 interactions with *Arabidopsis* homologs of OsGRF4, OsBES1, and OsIPA1 (Supplemental Fig. S9 and Supplemental Table S1) (Sun et al. 2023; Hu et al. 2020; Song et al. 2017).

Finally, we showed that chimeric SMXL proteins could be created that crosswire the normal responses to KARs and SLs. For example, the SMXLχ177 protein enabled SL-specific control of seed germination and seedling growth (Fig. 2). This suggests that important agronomic traits in crops that are controlled by SMXL proteins, including plant architecture and symbiotic interactions with microbes, could be genetically engineered to be regulated by a different hormone. We also demonstrated the creation of a miniaturized form of SMAX1 through the SMAX1_N210_-SRDX fusion. This protein recapitulates SMAX1 function but escapes SCF^MAX2^-dependent regulation (Fig. 3). It could conceivably be further fused to degrons from plant or non-plant systems to generate novel, inducible forms of developmental control (Huang and Rojas-Pierce 2024).

## METHODS

### Plant materials and growth conditions

*Arabidopsis thaliana* mutants *smax1-2 smxl2-1*, *max3-9*, and *max3-9 smxl6-4 smxl7-3 smxl8-1* were described previously (Stanga et al. 2016; Wang et al. 2015). Seeds were surface-sterilized, stratified at 4℃ for 3 d, plated on 0.5x Murashige and Skoog (0.5x MS) medium with 0.8% (w/v) Bacto agar, unless otherwise specified. For branching and height measurements, 10-d-old seedlings grown on 0.5xMS solid medium were transplanted to soil (Sungro Professional Growing Mix) supplemented with Gnatrol WDG and Marathon (imidacloprid) under 16-h white light/8-h dark cycles at ∼21℃.

### DNA constructs for transgenic plants

A binary Gateway destination vector, pGWBcitr, was generated by replacing the hygromycin resistance cassette of pGWB501 with a seed coat-specific Citrine cassette from pYUU (Nakagawa et al. 2007; Angulo et al. 2023). The promoter upstream of the Gateway cassette was replaced with *Arabidopsis SMAX1* and *SMXL7* promoter (3 kbp of DNA upstream of the translation start site). *Arabidopsis SMAX1* and *SMXL7* coding sequences with C-terminal 3xFLAG tags were cloned into pDONR221 vector by Gateway BP reaction (Invitrogen). SMXL chimeras were assembled by overlap-extension PCR (Nelson and Fitch 2011) and cloned into pDONR221. N-terminal SV40 NLS-, C-terminal FLAG-, and SRDX-fused *SMAX1_N_*, *SMAX1_N210_*, *SMXL7_N_, SMXL7_N190_* sequences were synthesized (Twist Biosciences) and cloned into pDONR221. *NLS-SMAX1_N_-FLAG, NLS- SMAX1_N210_-FLAG, NLS-SMXL7_N_-FLAG* and *NLS-SMXL7_N190_-FLAG* were amplified from the synthetic DNA by a FLAG specific primer and introduced into pDONR221 to generate Gateway entry clones. *NLS-eGFP-FLAG-SRDX* and *NLS-eGFP-FLAG* were assembled through NEBuilder (New England Biolabs) and cloned by BP reaction. Entry clones of *NLS-SMAX1ΔN-FLAG* and *NLS-SMXL7ΔN-FLAG* were generated with NEBuilder by replacing eGFP from an *NLS-eGFP-FLAG* entry clone with N-terminally truncated *SMAX1* and *SMXL7*. Entry clones of *SMAX1^ΔRGKT^*, *SMXL7^ΔRGKT^*, *SMXLχ177^ΔRGKT^*, and *SMXLχ1_210_77^ΔRGKT^* were generated with Q5 Site-Directed Mutagenesis Kit (New England Biolabs). Inserts in Gateway entry clones were transferred into pGWBcitr-*SMAX1pro* and pGWBcitr-*SMXL7pro* by Gateway LR reaction (Invitrogen). The eYFP-SMXLχ fusions used for testing subcellular localization were made by introducing the series of entry clones containing the intact and chimeric SMXLs into pGWB542 (Nakagawa et al. 2007) by Gateway LR reaction. Primers used for cloning, plasmid construction, and genotyping are listed in Supplemental Table S3.

### Structural analysis of SMAX1 and SMXL7

Globular regions of SMAX1 and SMXL7 were predicted with IUPred3 structural domains tool (Erdős et al. 2021). IDR regions were predicted using the D^2^P^2^ database (Oates et al. 2013). Predicted IDRs with over 75% agreement are indicated in Supplemental Fig. S7 and Supplemental Table S1.

### Plant phenotyping

Hypocotyl elongation assays were performed with slight modification as previously described (Sepulveda et al. 2022). Seeds were surface-sterilized, plated on 0.5xMS media supplemented with 1 µM KAR_2_, GR24^5DS^, or an equivalent volume of acetone solvent, stratified for 3 d at 4°C in darkness, and moved to a HiPoint DCI-700 LED Z4 growth chamber to grow at 21 °C under white light (150 µmol m^−2^ s^−1^) for 3 h, dark for 21 h, and continuous red light (30 µmol m^−2^ s^−1^) for 6 d. Hypocotyl lengths were measured from photographs of seedlings using ImageJ (NIH). Statistical significance (*P*<0.05) was calculated through Tukey’s multiple comparisons.

For branching and shoot height measurement, seedlings grown on 0.5xMS were moved to soil without fertilizer and grown as described above for 8 weeks. The primary shoot height and the number of rosette axillary branches at least 10 mm in length were counted. Statistical significance (*P*<0.05) was calculated through Tukey’s multiple comparisons. Germination assays of seeds aged at room temperature for at least one-month after harvest were performed as previously described (Bunsick et al. 2020). Seeds were surface-sterilized, plated on 0.5x MS containing 3 µM paclobutrazol (PAC), stratified 4 d at 4°C in darkness, and then moved to a HiPoint DCI-700 LED Z4 growth chamber to grow at 25°C under continuous white light (100 µmol m^−2^ s^−1^). After 10 d, germination was scored as radicle emergence.

### Yeast two-hybrid assays

*SMAX1, SMAX1ΔN, SMAX1_N_, SMAX1_N210_, SMXL7, SMXL7ΔN, SMXL7_N_,* and *SMXL7_N190_* in Gateway entry clones were transferred into pDEST32 (Invitrogen) by Gateway LR reaction. Bait plasmids were introduced into the Y2HGold yeast strain (Takara) using the lithium acetate method (Gietz and Schiestl 2007). Prey plasmids from the pDEST22- *Arabidopsis* TF library (Pruneda-Paz et al. 2014) were introduced into bait-transformed yeast lines. Co-transformed yeast were selected through growth on −Leu/−Trp (−LW) synthetic dropout media for 3 d at 30 °C. Bait-prey interactions were examined by spotting cells (10 μL of colony suspension at OD_600_ 0.15) on −LW and −Leu/−Trp/−His (−LWH) synthetic dropout plates. After 3 d at 30 °C, plates were photographed and colony growth was quantified using a modified density analysis method (Petropavlovskiy et al. 2020). Briefly, photographs were converted to grayscale, and gray values of yeast spots and their backgrounds were measured using ImageJ (NIH). Background values were subtracted from yeast spot values on the same plate. The background-subtracted values from −LWH plates were divided by the corresponding values from −LW plates. The ratios for the two colony replicates in each test were log_2_ transformed and averaged. Values above -0.5 were considered a positive interaction.

### Subcellular localization analysis

*A. tumefaciens* strain GV3101 carrying pGWB542 expression clones with *SMAX1*, *SMXL7*, and *SMXLχ* variants were infiltrated into *N. benthamiana* as previously described (Khosla et al. 2020b). The eYFP, 4′,6-diamino-2-phenylindole dihydrochloride (DAPI) and propidium iodide (PI) fluorescent signals were visualized using 880 Inverted Airyscan Fast confocal microscope (Zeiss) with the setting of eYFP (excitation, 514 nm; emission, 527 nm), DAPI (excitation, 405 nm; emission, 488 nm) and PI (excitation, 535/20 nm; emission, 610/20). For co-staining, leaf discs were incubated in distilled water supplemented with 10 µg/mL DAPI and 10 µg/mL PI for 20 min in darkness.

### Gene accession numbers

SMAX1 (AT5G57710.1), SMXL2 (AT4G30350.1), SMXL6 (AT1G07200.2), SMXL7 (AT2G29970.1), SMXL8 (AT2G40130.2), OsD53 (Os11g01330.1), OsSMAX1L (Os08g15230.1), OsSMXL2 (Os02g54720.1). Accession numbers for TFs used in the Y2H assay are listed in Supplemental Table S2.

## ACKNOWLEDGMENTS

Funding was provided by the National Science Foundation (NSF-IOS 1856741) and the United States Department of Agriculture (Hatch project CA-R-BPS-5209-H). We thank Dr. Bing Wang (Chinese Academy of Sciences, China) for providing *max3-9* and *max3-9 smxl6,7,8* lines. We thank Dr. Dawn H. Nagel (University of California, Riverside, USA) for sharing the *Arabidopsis* TF library.

## AUTHOR CONTRIBUTIONS

Conceptualization by S.C. and D.C.N. Investigation by S.C. and W.J.G. Formal analysis and visualization by S.C. Funding acquisition, project administration, and supervision by D.C.N. Writing by S.C. and D.C.N.

---

**Supplemental Figure 1.**
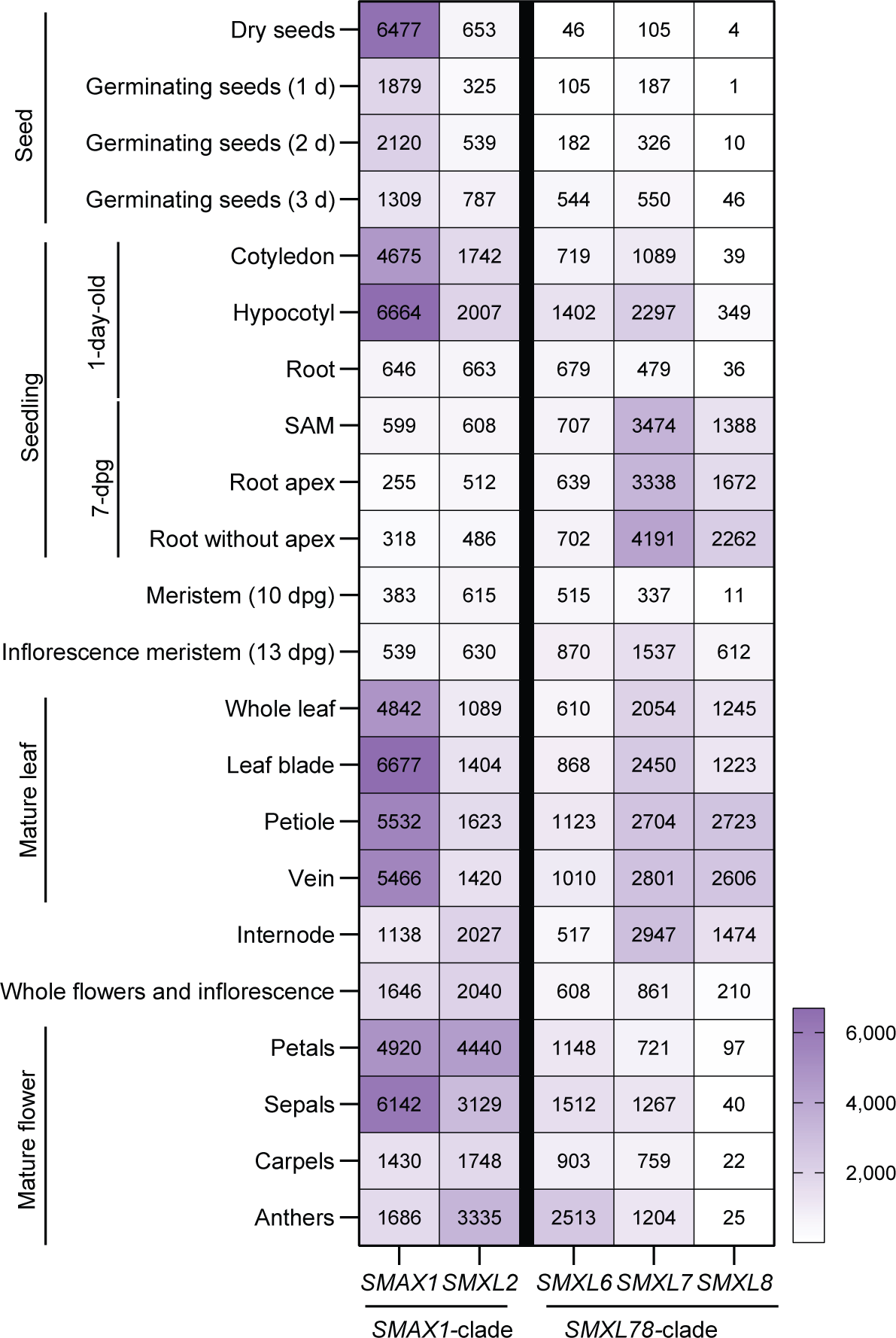
Expression pattern of *SMAX1*- and *SMXL78*-clade genes in different stages of *Arabidopsis* development. Overview of mRNA expression of *SMAX1* and *SMXL78* clades using TraVA (travadb.org) (Klepikova et al. 2016). The number in each box represents the normalized average count per million reads for each gene in the corresponding sample. ‘Seed’ samples were collected on the specified days after germination. Seedling samples labeled ‘1-day-old’ were collected from one-day-old seedlings, and ‘7 dpg’ samples were collected from the whole or specified tissues of the third leaf at the time of anthesis of the first flower. All the samples categorized in ‘Mature leaf’ were collected from the whole or specified tissue of the third leaf at the time of anthesis of the first flower. The ‘internode’ sample represents the first elongated internode between the last rosette leaf and the first cauline leaf, collected at the time of the anthesis of the first flower. ‘Whole flowers and inflorescence’ represents the average expression of flowers collected at the time of anthesis of the first flower. Samples of ‘Mature flower’ were collected from the specified floral parts collected at the moment of the anthesis of the first flower. ‘D’ means day, and ‘dpg’ means day post-germination. More detailed information on the samples can be found in http://travadb.org/samples/.

**Supplemental Figure 2.**
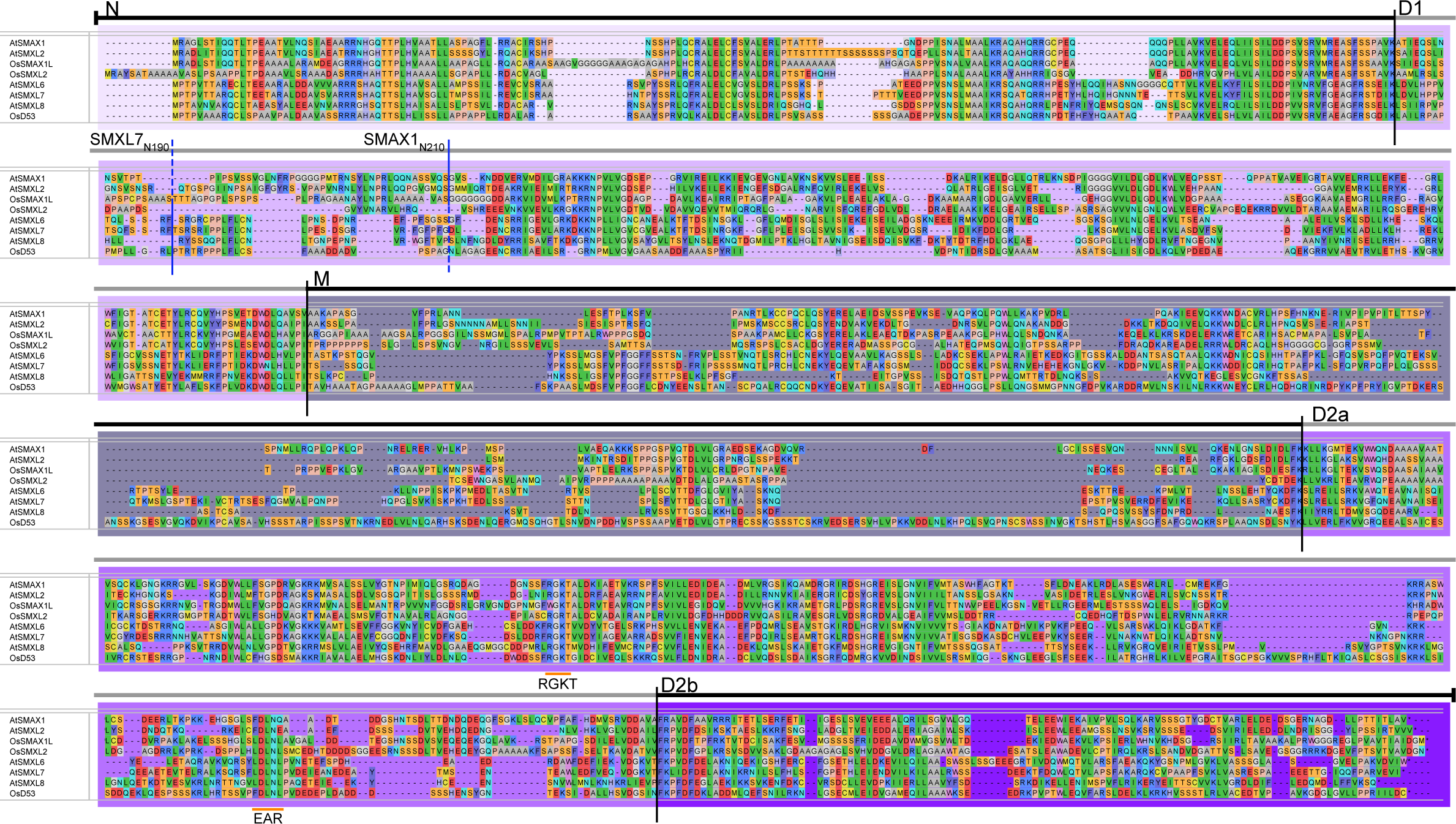
Domain boundaries of SMAX1- and SMXL78-clade proteins in *Arabidopsis* and rice. Multiple sequence alignment of SMAX1 and SMXL78-clade protein sequences from *Arabidopsis thaliana* and *Oryza sativa* (rice) to show the domain boundaries. The start positions of the N, D1, M, D2a, and D2b domains are indicated by black vertical bars and each domain is color-labeled with purple. The extended boundaries for SMAX1_N210_ and SMXL7_N190_ are marked with blue bars. RGKT motif and EAR motif are orange highlighted. The residues are highlighted based on their chemical properties. The alignment was performed using Clustal Omega in DNASTAR MegAlign software.

**Supplemental Figure 3.**
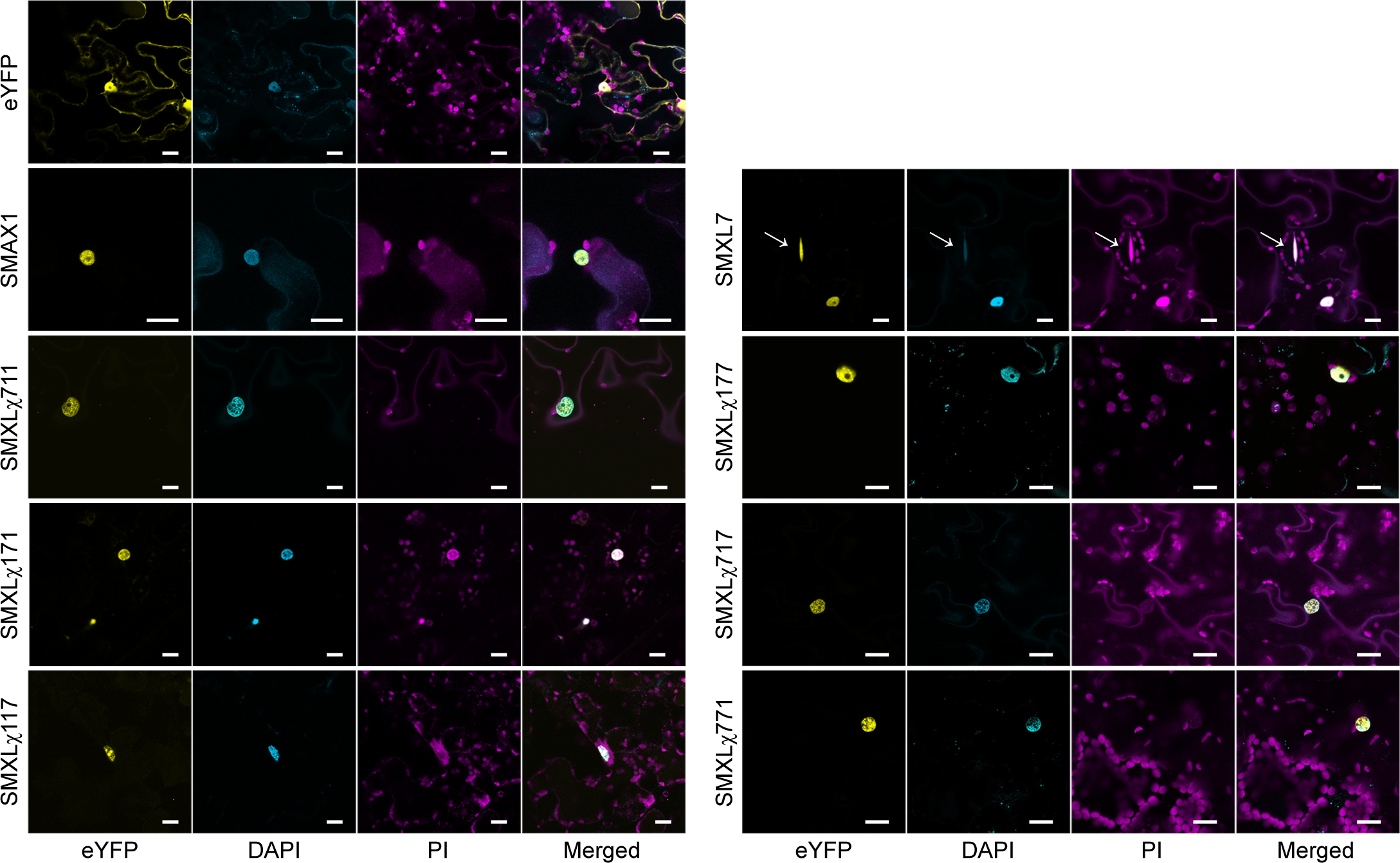
Subcellular localization of eYFP-tagged chimeric SMXLs in *N. benthamiana*. Confocal microscopy images of transiently expressed N-terminal eYFP-tagged SMAX1, SMXL7, and chimeric SMXLs in *N. benthamiana*. The leaf discs were stained with DAPI and PI. Arrows indicate stomata. Scale bar = 20 µm.

**Supplemental Figure 4.**
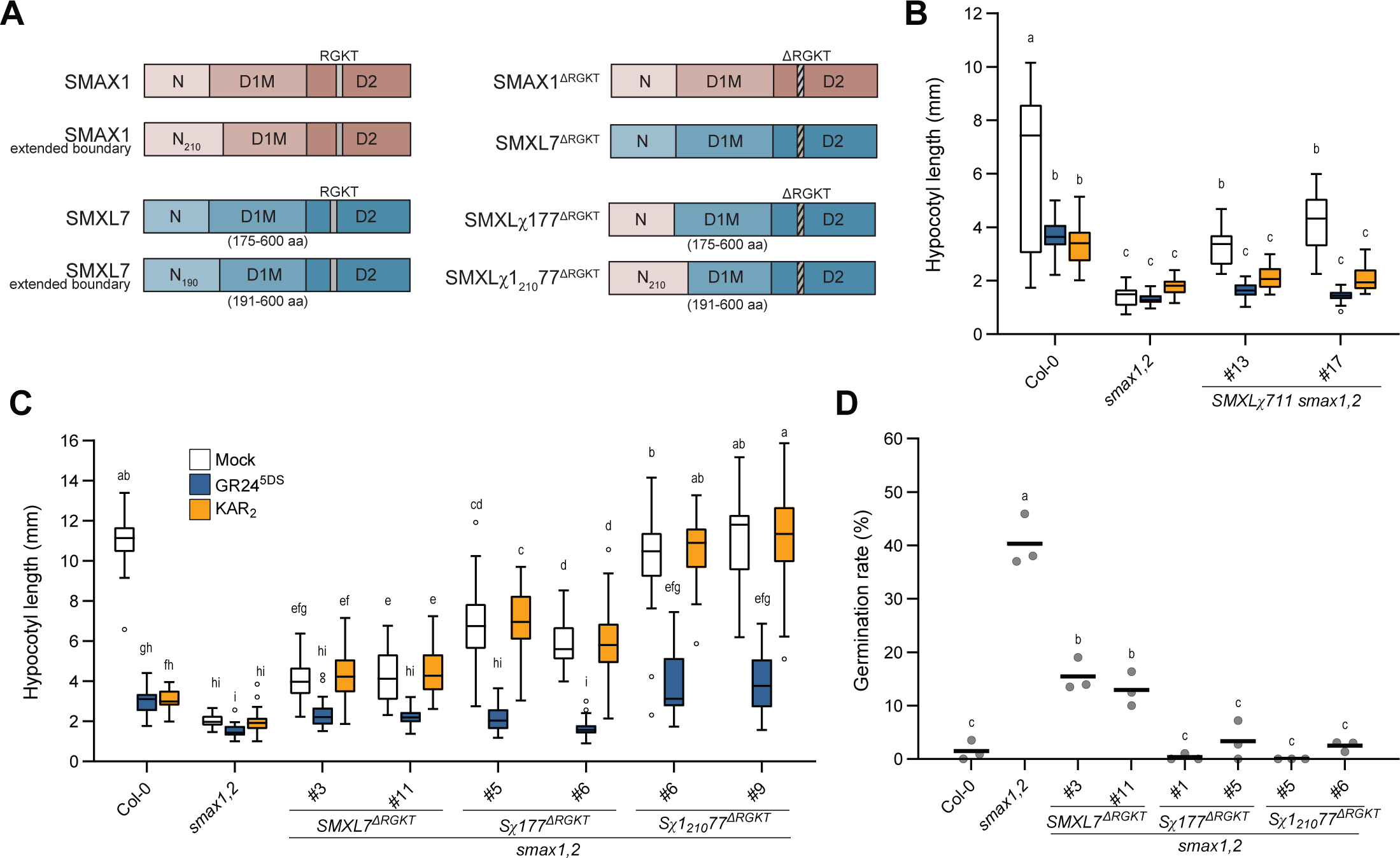
Seed and seedling growth control by SMXL N domains. (*A*) Diagram of SMAX1, SMXL7, SMXL!177, SMXL!1_210_77, and their RGKT motif-deleted versions SMAX1^ΔRGKT^, SMXL7^ΔRGKT^, SMXL!177^ΔRGKT^, and SMXL!1_210_77^ΔRGKT^. The positions of the RGKT motif and the extended N domain boundaries are indicated. (*B*) Hypocotyl length of Col-0, *smax1,2* and two independent lines of *SMAX1pro::SMXL!177 smax1,2*. Seedlings were treated with mock (acetone), 1 μM KAR_2_, or 1 μM GR24^5DS^ (n>20). (*C*) Hypocotyl length of Col-0, *smax1,2*, and transgenic lines expressing *SMXL7^ΔRGKT^*, *SMXL!177^ΔRGKT^*, and *SMXL!1_210_77^ΔRGKT^* under control of the *SMXL7* promoter with mock, 1 μM KAR_2_, or 1 μM GR24^5DS^ treatment (n>20). (*D*) Germination of the lines in *C* (n=3, >50 seeds per replicate). Boxplots indicate mean with quartiles and Tukey’s whiskers; open symbols are outlier points that fall beyond the range of the whiskers. Letters indicate groups with significant differences (*P*<0.05, two-way ANOVA in *B* and *C,* or one-way ANOVA in *D*, followed by Tukey’s multiple comparisons test). S!, abbreviation for SMXL!.

**Supplemental Figure 5.**
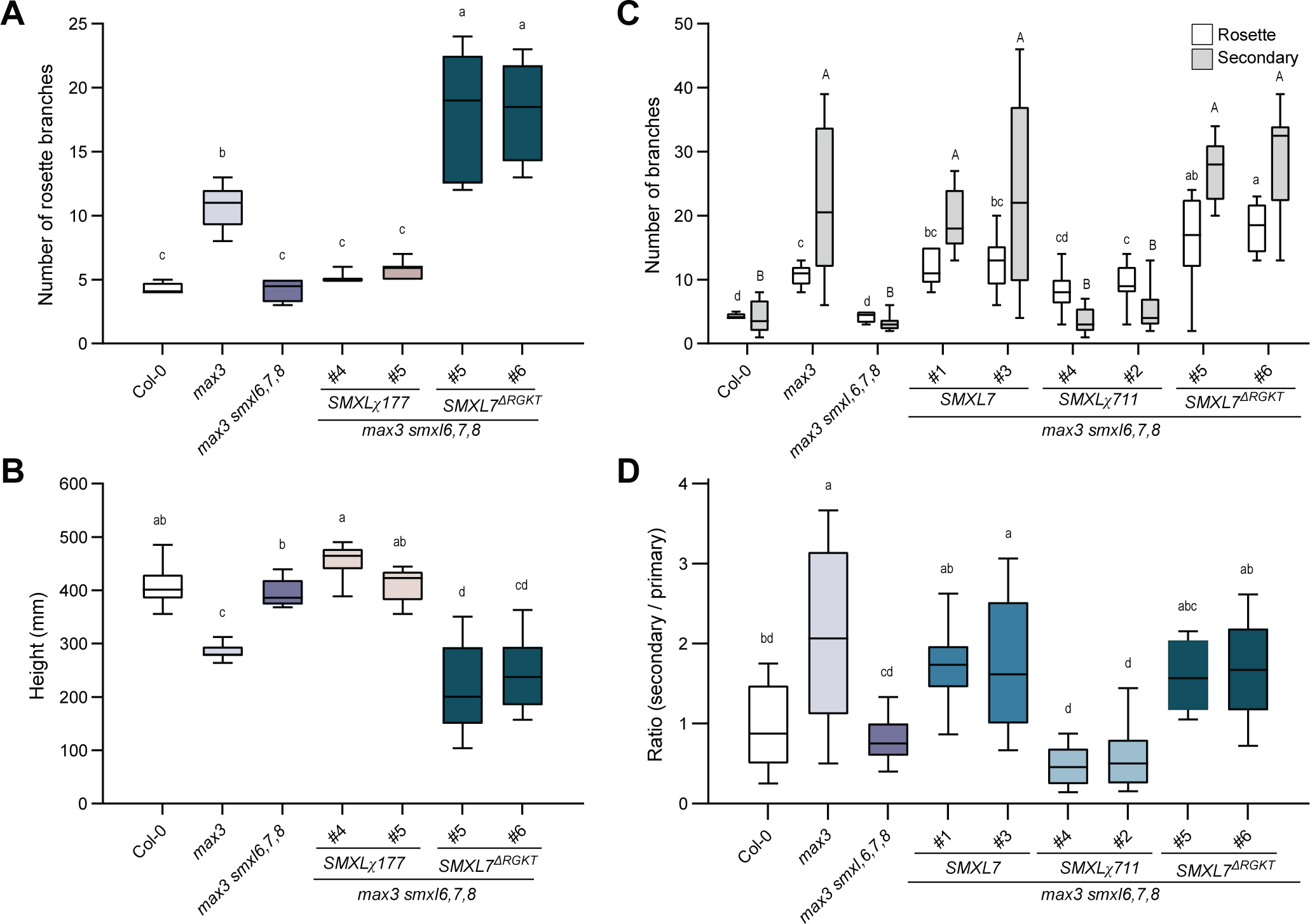
SMXL7N is not sufficient to rescue *max3 smxl6,7,8*. (A) Number of axillary branches in transgenic lines expressing *SMXL7pro::SMXL!177* and *SMXL7pro::SMXL7^ΔRGKT^* in *max3 smxl6,7,8* (n>9). Data for the Col-0, *max3,* and *max3 smxl6,7,8* control plants is duplicated in Figure 2. (*B*) Plant height of the lines used in *A*. (*C*) Number of rosette branches (white) and secondary branches (gray) of the indicated transgenics. Data for the *SMXL7pro::SMXL7* and *SMXL7pro::SMXL!711* lines is duplicated in Figure 2D, and data for the *SMXL7pro::SMXL7^ΔRGKT^* lines is duplicated in *A-B*. (*D*) Ratio of secondary branches to rosette branches in the lines used in *C*. Boxplots indicate mean with quartiles and Tukey’s whiskers; open symbols are outlier points that fall beyond the range of the whiskers. Letters indicate groups with significant differences (*P*<0.05, one-way ANOVA followed by Tukey’s multiple comparisons test). In *C*, significant differences were calculated separately for rosette branches (uppercase letters) and secondary branches (lowercase letters).

**Supplemental Figure 6.**
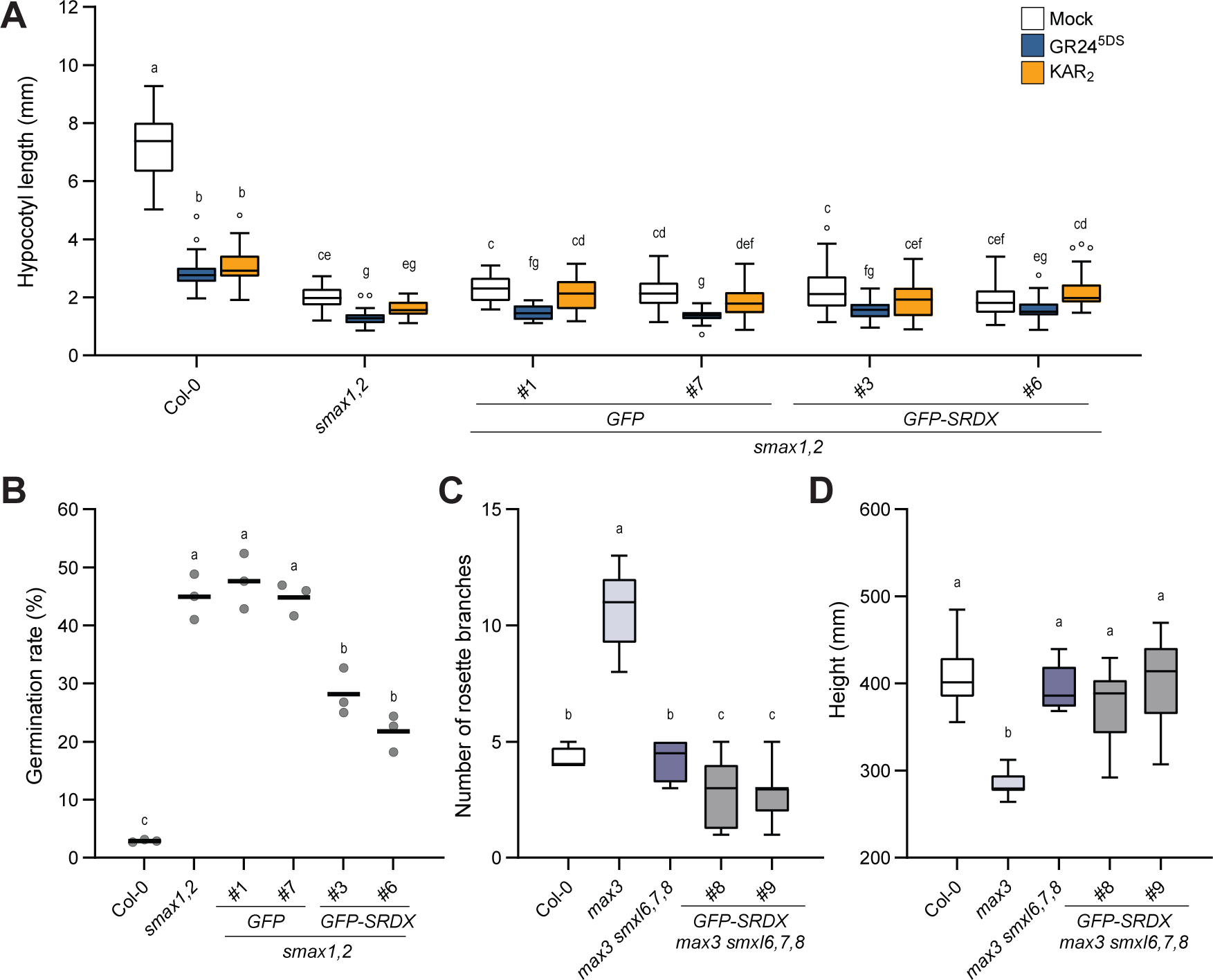
GFP-SRDX does not rescue *smax1,2* or *max3 smxl6,7,8*. (*A*) Hypocotyl length of Col-0, *smax1,2* and two independent transgenic lines expressing *SMAX1pro::GFP* or *SMAX1pro::GFP-SRDX* in *smax1,2.* Seedlings were treated with mock (acetone), 1 μM KAR_2_, or 1 μM GR24^5DS^ (n>20). These lines were tested together with those in Figure 3B; Col-0 and *smax1,2* data is duplicated in Figure 3B. (*B*) Germination rate of the lines used in a (n=3, >50 seeds per replicate). (*C*) Number of rosette axillary branches in transgenic lines expressing *SMXL7pro::GFP-SRDX* in *max3 smxl6,7,8* (n>9). Data for the Col-0, *max3,* and *max3 smxl6,7,8* plants is duplicated in Figure 2. (*D*) Plant height of the lines used in *C*. Letters indicate groups with significant differences (*P*<0.05, two-way ANOVA in *A*, or one-way ANOVA in *B-D*, followed by Tukey’s multiple comparisons test).

**Supplemental Figure 7.**
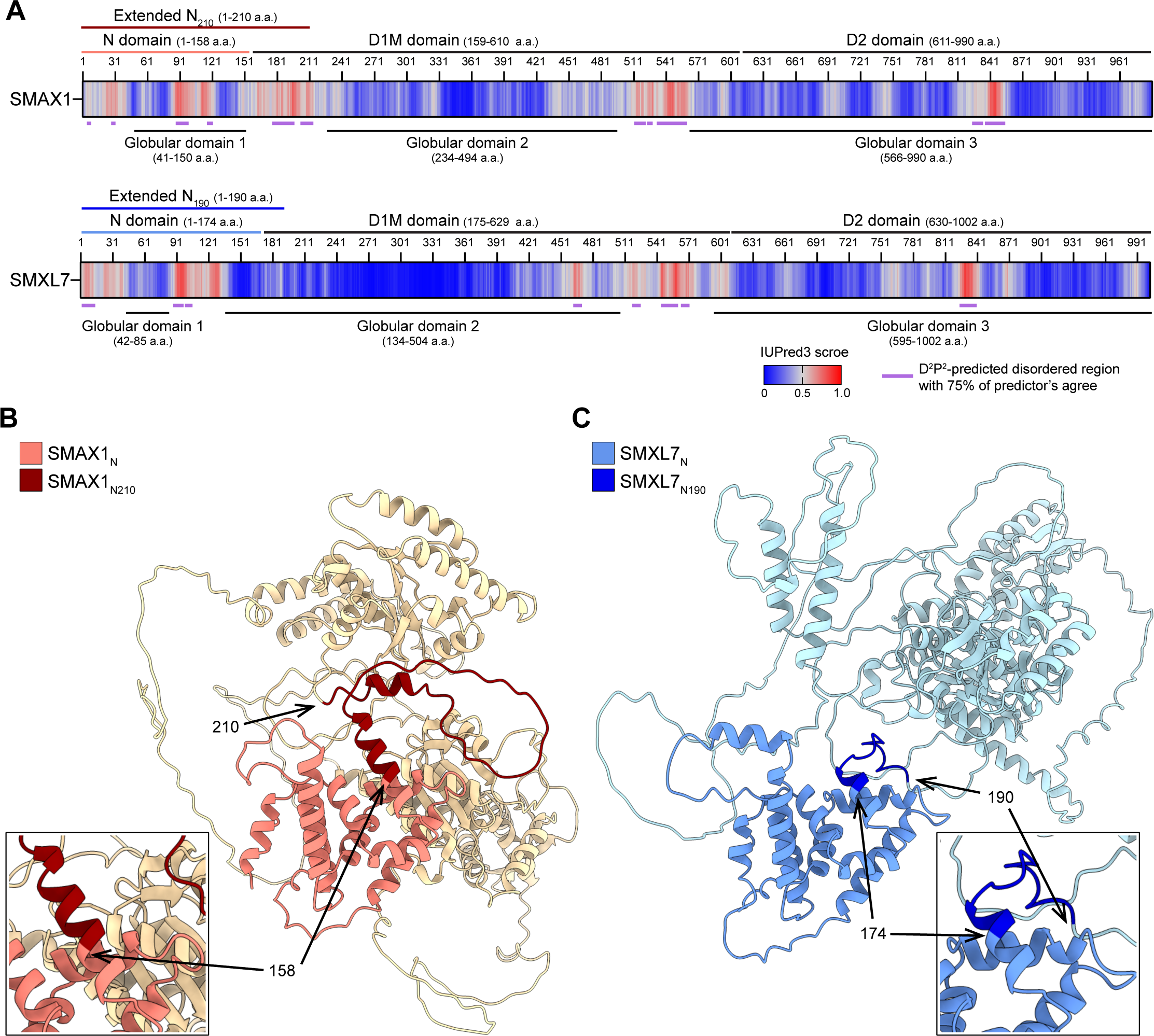
The boundaries of the defined N domains for SMAX1 and SMXL7 do not include the entirety of the last predicted alpha helix in the domain. (*A*) The prediction of globular domains and IDRs of SMAX1 and SMXL7. Globular domains were predicted using IUPred3, and disordered regions were predicted using D^2^P^2^. D^2^P^2^-predicted IDRs longer than 4 amino acids are represented by purple bars in the figure. Their detailed positions are listed in Supplemental Table 1. (*B*) The Alphafold2-predicted structure of SMAX1 and SMXL7. The N domains of SMAX1 and SMXL7 are highlighted in salmon pink and blue, respectively. The extended SMAX1 N domain (SMAX1_N210_) and SMXL7 N domain (SMXL7_N190_) are colored dark red and dark blue. The alpha helix regions at the N domain boundaries are magnified in the inset boxes.

**Supplemental Figure 8.**
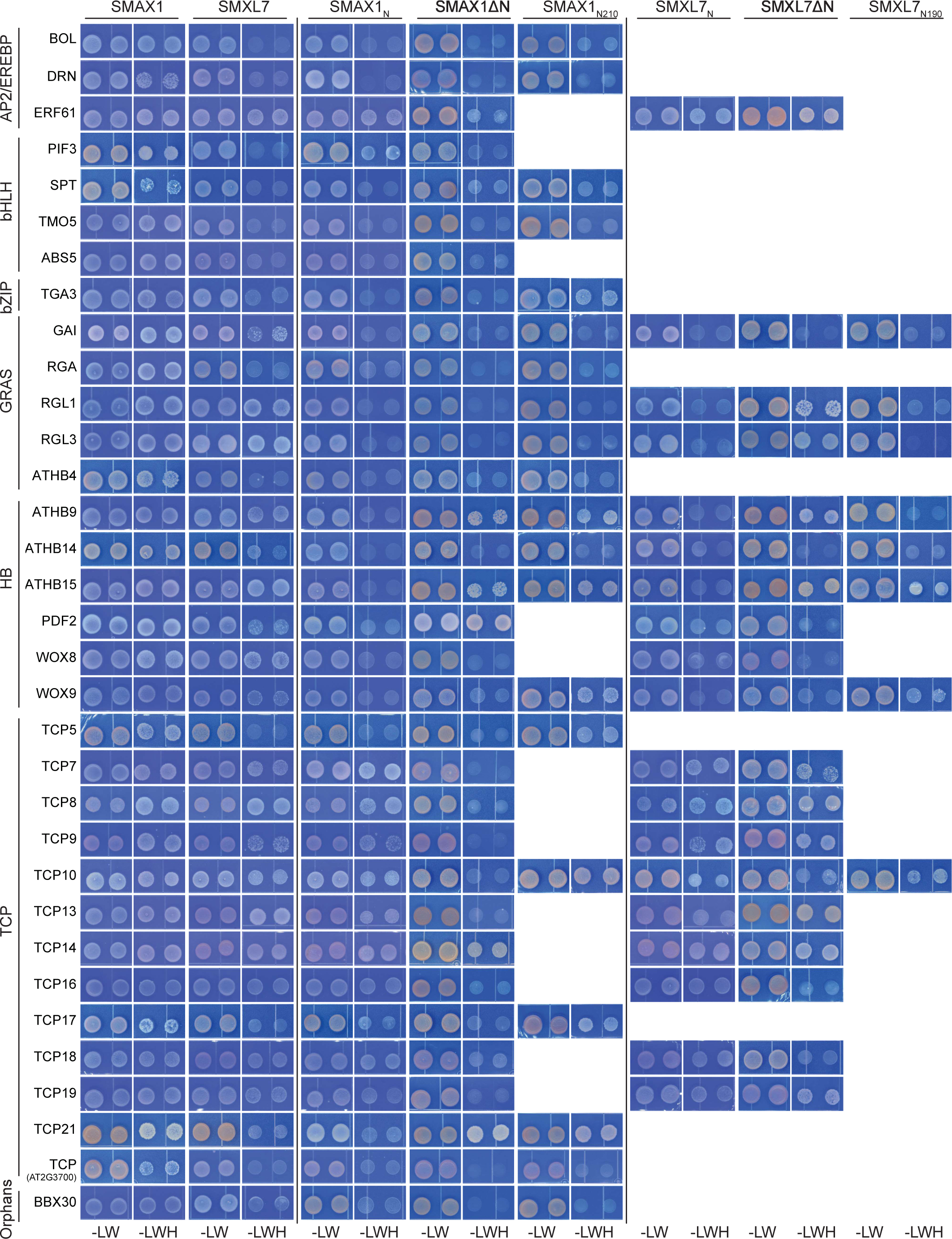
Photograph of Y2H results with TFs showing potential interactions with SMAX1 and/or SMXL7. Interactions between SMAX1, SMXL7 and their truncated variants fused to GAL4-BD and candidate TFs fused to GAL4-AD were tested. Two replicates were spotted onto selective growth medium (-L, -Leu; -W, -Trp; -H, - His), incubated 3 d at 30℃, and photographed.

**Supplemental Figure 9.**
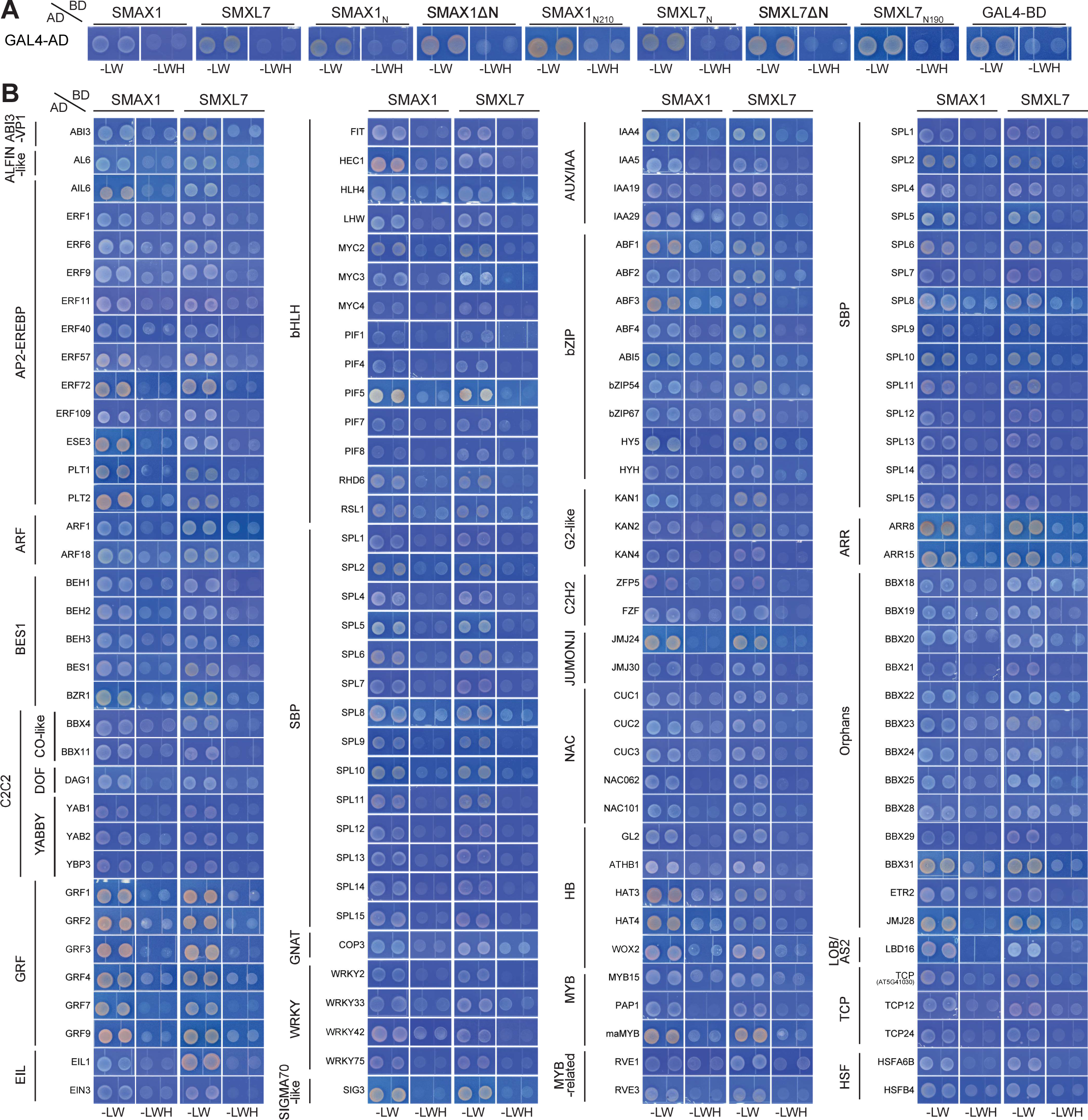
Y2H assays that showed no interaction between SMAX1 and candidate TFs. (*A*) Autoactivation test for SMAX1, SMXL7, and their truncated versions fused with the GAL4-BD. (*B*) Y2H assay results showing no interaction between SMAX1 or SMXL7 and the indicated TFs fused to GAL4-AD.

## REFERENCES

Aguilar-Martínez JA, Poza-Carrión C, Cubas P. 2007. Arabidopsis BRANCHED1 acts as an integrator of branching signals within axillary buds. Plant Cell 19: 458–472. 10.1105/tpc.106.048934.

Agusti J, Herold S, Schwarz M, Sanchez P, Ljung K, Dun EA, Brewer PB, Beveridge CA, Sieberer T, Sehr EM, et al. 2011. Strigolactone signaling is required for auxin-dependent stimulation of secondary growth in plants. Proc Natl Acad Sci U S A 108: 20242–20247. 10.1073/pnas.1111902108.

Angulo J, Astin CP, Bauer O, Blash KJ, Bowen NM, Chukwudinma NJ, DiNofrio AS, Faletti DO, Ghulam AM, Gusinde-Duffy CM, et al. 2023. CRISPR/Cas9 mutagenesis of the GROWTH-REGULATING FACTOR (GRF) gene family. Front Genome Ed 5: 1251557. 10.3389/fgeed.2023.1251557.

Blázquez MA, Nelson DC, Weijers D. 2020. Evolution of Plant Hormone Response Pathways. Annu Rev Plant Biol 71: 327–353. 10.1146/annurev-arplant-050718-100309.

Booker J, Auldridge M, Wills S, McCarty D, Klee H, Leyser O. 2004. MAX3/CCD7 is a carotenoid cleavage dioxygenase required for the synthesis of a novel plant signaling molecule. Curr Biol 14: 1232–1238. 10.1016/j.cub.2004.06.061.

Bunsick M, Toh S, Wong C, Xu Z, Ly G, McErlean CSP, Pescetto G, Nemrish KE, Sung P, Li JD, et al. 2020. SMAX1-dependent seed germination bypasses GA signalling in Arabidopsis and Striga. Nat Plants 6: 646–652. 10.1038/s41477-020-0653-z.

Bursch K, Niemann ET, Nelson DC, Johansson H. 2021. Karrikins control seedling photomorphogenesis and anthocyanin biosynthesis through a HY5-BBX transcriptional module. Plant J 107: 1346–1362. 10.1111/tpj.15383.

Carbonnel S, Das D, Varshney K, Kolodziej MC, Villaécija-Aguilar JA, Gutjahr C. 2020a. The karrikin signaling regulator SMAX1 controls Lotus japonicus root and root hair development by suppressing ethylene biosynthesis. Proc Natl Acad Sci U S A 117: 21757–21765. 10.1073/pnas.2006111117.

Carbonnel S, Torabi S, Griesmann M, Bleek E, Tang Y, Buchka S, Basso V, Shindo M, Boyer F- D, Wang TL, et al. 2020b. Lotus japonicus karrikin receptors display divergent ligand-binding specificities and organ-dependent redundancy. PLoS Genet 16: e1009249. 10.1371/journal.pgen.1009249.

Cho H, Cho HS, Nam H, Jo H, Yoon J, Park C, Dang TVT, Kim E, Jeong J, Park S, et al. 2018. Translational control of phloem development by RNA G-quadruplex-JULGI determines plant sink strength. Nat Plants 4: 376–390. 10.1038/s41477-018-0157-2.

Choi J, Lee T, Cho J, Servante EK, Pucker B, Summers W, Bowden S, Rahimi M, An K, An G, et al. 2020. The negative regulator SMAX1 controls mycorrhizal symbiosis and strigolactone biosynthesis in rice. Nat Commun 11: 2114. 10.1038/s41467-020-16021-1.

Conn CE, Bythell-Douglas R, Neumann D, Yoshida S, Whittington B, Westwood JH, Shirasu K, Bond CS, Dyer KA, Nelson DC. 2015. PLANT EVOLUTION. Convergent evolution of strigolactone perception enabled host detection in parasitic plants. Science 349: 540–543. 10.1126/science.aab1140.

Conn CE, Nelson DC. 2015. Evidence that KARRIKIN-INSENSITIVE2 (KAI2) Receptors may Perceive an Unknown Signal that is not Karrikin or Strigolactone. Front Plant Sci 6: 1219. 10.3389/fpls.2015.01219.

de Saint Germain A, Clavé G, Badet-Denisot M-A, Pillot J-P, Cornu D, Le Caer J-P, Burger M, Pelissier F, Retailleau P, Turnbull C, et al. 2016. An histidine covalent receptor and butenolide complex mediates strigolactone perception. Nat Chem Biol 12: 787–794. 10.1038/nchembio.2147.

de Saint Germain A, Jacobs A, Brun G, Pouvreau J-B, Braem L, Cornu D, Clavé G, Baudu E, Steinmetz V, Servajean V, et al. 2021. A Phelipanche ramosa KAI2 protein perceives strigolactones and isothiocyanates enzymatically. Plant Communications 2: 100166. https://www.sciencedirect.com/science/article/pii/S2590346221000419.

de Saint Germain A, Ligerot Y, Dun EA, Pillot J-P, Ross JJ, Beveridge CA, Rameau C. 2013. Strigolactones stimulate internode elongation independently of gibberellins. Plant Physiol 163: 1012–1025. 10.1104/pp.113.220541.

Erdős G, Pajkos M, Dosztányi Z. 2021. IUPred3: prediction of protein disorder enhanced with unambiguous experimental annotation and visualization of evolutionary conservation. Nucleic Acids Res 49: W297–W303. 10.1093/nar/gkab408.

Fang Z, Ji Y, Hu J, Guo R, Sun S, Wang X. 2020. Strigolactones and Brassinosteroids Antagonistically Regulate the Stability of the D53-OsBZR1 Complex to Determine FC1 Expression in Rice Tillering. Mol Plant 13: 586–597. 10.1016/j.molp.2019.12.005.

Feng Z, Liang X, Tian H, Watanabe Y, Nguyen KH, Tran CD, Abdelrahman M, Xu K, Mostofa MG, Van Ha C, et al. 2022. SUPPRESSOR of MAX2 1 (SMAX1) and SMAX1-LIKE2 (SMXL2) Negatively Regulate Drought Resistance in Arabidopsis thaliana. Plant Cell Physiol. 10.1093/pcp/pcac080.

Gallie DR, Fortner D, Peng J, Puthoff D. 2002. ATP-dependent hexameric assembly of the heat shock protein Hsp101 involves multiple interaction domains and a functional C-proximal nucleotide-binding domain. J Biol Chem 277: 39617–39626. 10.1074/jbc.M204998200.

Gietz RD, Schiestl RH. 2007. Frozen competent yeast cells that can be transformed with high efficiency using the LiAc/SS carrier DNA/PEG method. Nat Protoc 2: 1–4. 10.1038/nprot.2007.17.

Guercio AM, Torabi S, Cornu D, Dalmais M, Bendahmane A, Le Signor C, Pillot J-P, Le Bris P, Boyer F-D, Rameau C, et al. 2022. Structural and functional analyses explain Pea KAI2 receptor diversity and reveal stereoselective catalysis during signal perception. Commun Biol 5: 126. 10.1038/s42003-022-03085-6.

Hamiaux C, Drummond RSM, Janssen BJ, Ledger SE, Cooney JM, Newcomb RD, Snowden KC. 2012. DAD2 is an α/β hydrolase likely to be involved in the perception of the plant branching hormone, strigolactone. Curr Biol 22: 2032–2036. 10.1016/j.cub.2012.08.007.

Hiratsu K, Matsui K, Koyama T, Ohme-Takagi M. 2003. Dominant repression of target genes by chimeric repressors that include the EAR motif, a repression domain, in Arabidopsis. Plant J 34: 733–739. 10.1046/j.1365-313x.2003.01759.x.

Huang L, Rojas-Pierce M. 2024. Rapid depletion of target proteins in plants by an inducible protein degradation system. Plant Cell. 10.1093/plcell/koae072.

Hu J, Ji Y, Hu X, Sun S, Wang X. 2020. BES1 Functions as the Co-regulator of D53-like SMXLs to Inhibit BRC1 Expression in Strigolactone-Regulated Shoot Branching in Arabidopsis. Plant Commun 1: 100014. 10.1016/j.xplc.2019.100014.

Humphreys JL, Beveridge C, Tanurdzic M. 2023. Strigolactone-dependent gene regulation requires chromatin remodeling. bioRxiv 2023.02.25.529999. https://www.biorxiv.org/content/10.1101/2023.02.25.529999v1 (Accessed April 18, 2024).

Jiang L, Liu X, Xiong G, Liu H, Chen F, Wang L, Meng X, Liu G, Yu H, Yuan Y, et al. 2013. DWARF 53 acts as a repressor of strigolactone signalling in rice. Nature 504: 401–405. 10.1038/nature12870.

Jumper J, Evans R, Pritzel A, Green T, Figurnov M, Ronneberger O, Tunyasuvunakool K, Bates R, Žídek A, Potapenko A, et al. 2021. Highly accurate protein structure prediction with AlphaFold. Nature 596: 583–589. 10.1038/s41586-021-03819-2.

Khosla A, Morffy N, Li Q, Faure L, Chang SH, Yao J, Zheng J, Cai ML, Stanga J, Flematti GR, et al. 2020a. Structure-Function Analysis of SMAX1 Reveals Domains That Mediate Its Karrikin-Induced Proteolysis and Interaction with the Receptor KAI2. Plant Cell 32: 2639–2659. 10.1105/tpc.19.00752.

Khosla A, Rodriguez-Furlan C, Kapoor S, Van Norman JM, Nelson DC. 2020b. A series of dual-reporter vectors for ratiometric analysis of protein abundance in plants. Plant Direct 4: e00231. 10.1002/pld3.231.

Kim JY, Park Y-J, Lee J-H, Park C-M. 2022. SMAX1 Integrates Karrikin and Light Signals into GA-Mediated Hypocotyl Growth during Seedling Establishment. Plant Cell Physiol 63: 932– 943. 10.1093/pcp/pcac055.

Klepikova AV, Kasianov AS, Gerasimov ES, Logacheva MD, Penin AA. 2016. A high resolution map of the Arabidopsis thaliana developmental transcriptome based on RNA-seq profiling. Plant J 88: 1058–1070. 10.1111/tpj.13312.

Liang Y, Ward S, Li P, Bennett T, Leyser O. 2016. SMAX1-LIKE7 Signals from the Nucleus to Regulate Shoot Development in Arabidopsis via Partially EAR Motif-Independent Mechanisms. Plant Cell 28: 1581–1601. 10.1105/tpc.16.00286.

Li Q, Martín-Fontecha ES, Khosla A, White ARF, Chang S, Cubas P, Nelson DC. 2022. The strigolactone receptor D14 targets SMAX1 for degradation in response to GR24 treatment and osmotic stress. Plant Communications 100303. https://www.sciencedirect.com/science/article/pii/S2590346222000505.

Li Q, Yu H, Chang W, Chang S, Guzmán M, Faure L, Wallner E-S, Yan H, Greb T, Wang L, et al. 2024. SMXL5 attenuates strigolactone signaling in Arabidopsis thaliana by inhibiting SMXL7 degradation. Mol Plant 17: 631–647. 10.1016/j.molp.2024.03.006.

Li W, Nguyen KH, Tran CD, Watanabe Y, Tian C, Yin X, Li K, Yang Y, Guo J, Miao Y, et al. 2020. Negative Roles of Strigolactone-Related SMXL6, 7 and 8 Proteins in Drought Resistance in Arabidopsis. Biomolecules 10. 10.3390/biom10040607.

Ma H, Duan J, Ke J, He Y, Gu X, Xu T-H, Yu H, Wang Y, Brunzelle JS, Jiang Y, et al. 2017. A D53 repression motif induces oligomerization of TOPLESS corepressors and promotes assembly of a corepressor-nucleosome complex. Sci Adv 3: e1601217. 10.1126/sciadv.1601217.

Martinez SE, Conn CE, Guercio AM, Sepulveda C, Fiscus CJ, Koenig D, Shabek N, Nelson DC. 2022. A KARRIKIN INSENSITIVE2 paralog in lettuce mediates highly sensitive germination responses to karrikinolide. Plant Physiol 190: 1440–1456. 10.1093/plphys/kiac328 (Accessed March 29, 2022).

Nakagawa T, Suzuki T, Murata S, Nakamura S, Hino T, Maeo K, Tabata R, Kawai T, Tanaka K, Niwa Y, et al. 2007. Improved Gateway binary vectors: high-performance vectors for creation of fusion constructs in transgenic analysis of plants. Biosci Biotechnol Biochem 71: 2095– 2100. 10.1271/bbb.70216.

Nelson MD, Fitch DHA. 2011. Overlap Extension PCR: An Efficient Method for Transgene Construction. In Molecular Methods for Evolutionary Genetics (eds. V. Orgogozo and M.V. Rockman), pp. 459–470, Humana Press, Totowa, NJ 10.1007/978-1-61779-228-1_27.

Oates ME, Romero P, Ishida T, Ghalwash M, Mizianty MJ, Xue B, Dosztányi Z, Uversky VN, Obradovic Z, Kurgan L, et al. 2013. D^2^P^2^: database of disordered protein predictions. Nucleic Acids Res 41: D508–16. 10.1093/nar/gks1226.

Park Y-J, Kim JY, Park C-M. 2022. SMAX1 potentiates phytochrome B-mediated hypocotyl thermomorphogenesis. Plant Cell 34: 2671–2687. 10.1093/plcell/koac124.

Petropavlovskiy AA, Tauro MG, Lajoie P, Duennwald ML. 2020. A Quantitative Imaging-Based Protocol for Yeast Growth and Survival on Agar Plates. STAR Protoc 1: 100182. 10.1016/j.xpro.2020.100182.

Pruneda-Paz JL, Breton G, Nagel DH, Kang SE, Bonaldi K, Doherty CJ, Ravelo S, Galli M, Ecker JR, Kay SA. 2014. A genome-scale resource for the functional characterization of Arabidopsis transcription factors. Cell Rep 8: 622–632. 10.1016/j.celrep.2014.06.033.

Sepulveda C, Guzmán MA, Li Q, Villaécija-Aguilar JA, Martinez SE, Kamran M, Khosla A, Liu W, Gendron JM, Gutjahr C, et al. 2022. KARRIKIN UP-REGULATED F-BOX 1 (KUF1) imposes negative feedback regulation of karrikin and KAI2 ligand metabolism in Arabidopsis thaliana. Proc Natl Acad Sci U S A 119: e2112820119. 10.1073/pnas.2112820119.

Song X, Lu Z, Yu H, Shao G, Xiong J, Meng X, Jing Y, Liu G, Xiong G, Duan J, et al. 2017. IPA1 functions as a downstream transcription factor repressed by D53 in strigolactone signaling in rice. Cell Res 27: 1128–1141. 10.1038/cr.2017.102.

Soundappan I, Bennett T, Morffy N, Liang Y, Stanga JP, Abbas A, Leyser O, Nelson DC. 2015. SMAX1-LIKE/D53 Family Members Enable Distinct MAX2-Dependent Responses to Strigolactones and Karrikins in Arabidopsis. Plant Cell 27: 3143–3159. 10.1105/tpc.15.00562.

Stanga JP, Morffy N, Nelson DC. 2016. Functional redundancy in the control of seedling growth by the karrikin signaling pathway. Planta 243: 1397–1406. 10.1007/s00425-015-2458-2.

Stanga JP, Smith SM, Briggs WR, Nelson DC. 2013. SUPPRESSOR OF MORE AXILLARY GROWTH2 1 controls seed germination and seedling development in Arabidopsis. Plant Physiol 163: 318–330. 10.1104/pp.113.221259.

Stirling SA, Guercio AM, Patrick RM, Huang X-Q, Bergman ME, Dwivedi V, Kortbeek RWJ, Liu Y-K, Sun F, Tao WA, et al. 2024. Volatile communication in plants relies on a KAI2-mediated signaling pathway. Science 383: 1318–1325. 10.1126/science.adl4685.

Sun H, Guo X, Zhu X, Gu P, Zhang W, Tao W, Wang D, Wu Y, Zhao Q, Xu G, et al. 2023. Strigolactone and gibberellin signaling coordinately regulate metabolic adaptations to changes in nitrogen availability in rice. Mol Plant 16: 588–598. 10.1016/j.molp.2023.01.009.

Sun YK, Yao J, Scaffidi A, Melville KT, Davies SF, Bond CS, Smith SM, Flematti GR, Waters MT. 2020. Divergent receptor proteins confer responses to different karrikins in two ephemeral weeds. Nat Commun 11: 1264. 10.1038/s41467-020-14991-w.

Tal L, Palayam M, Ron M, Young A, Britt A, Shabek N. 2022. A conformational switch in the SCF-D3/MAX2 ubiquitin ligase facilitates strigolactone signalling. Nat Plants 8: 561–573. 10.1038/s41477-022-01145-7.

Temmerman A, Guillory A, Bonhomme S, Goormachtig S, Struk S. 2022. Masks Start to Drop: Suppressor of MAX2 1-Like Proteins Reveal Their Many Faces. Front Plant Sci 13: 887232. 10.3389/fpls.2022.887232.

Toh S, Holbrook-Smith D, Stogios PJ, Onopriyenko O, Lumba S, Tsuchiya Y, Savchenko A, McCourt P. 2015. Structure-function analysis identifies highly sensitive strigolactone receptors in Striga. Science 350: 203–207. 10.1126/science.aac9476.

Tsuchiya Y, Yoshimura M, Sato Y, Kuwata K, Toh S, Holbrook-Smith D, Zhang H, McCourt P, Itami K, Kinoshita T, et al. 2015. PARASITIC PLANTS. Probing strigolactone receptors in Striga hermonthica with fluorescence. Science 349: 864–868. 10.1126/science.aab3831.

Villaécija-Aguilar JA, Körösy C, Maisch L, Hamon-Josse M, Petrich A, Magosch S, Chapman P, Bennett T, Gutjahr C. 2022. KAI2 promotes Arabidopsis root hair elongation at low external phosphate by controlling local accumulation of AUX1 and PIN2. Curr Biol 32: 228–236.e3. 10.1016/j.cub.2021.10.044.

Walker CH, Siu-Ting K, Taylor A, O’Connell MJ, Bennett T. 2019. Strigolactone synthesis is ancestral in land plants, but canonical strigolactone signalling is a flowering plant innovation. BMC Biol 17: 70. 10.1186/s12915-019-0689-6.

Wallner E-S, López-Salmerón V, Belevich I, Poschet G, Jung I, Grünwald K, Sevilem I, Jokitalo E, Hell R, Helariutta Y, et al. 2017. Strigolactone- and Karrikin-Independent SMXL Proteins Are Central Regulators of Phloem Formation. Curr Biol 27: 1241–1247. 10.1016/j.cub.2017.03.014.

Wallner E-S, Tonn N, Shi D, Luzzietti L, Wanke F, Hunziker P, Xu Y, Jung I, Lopéz-Salmerón V, Gebert M, et al. 2023. OBERON3 and SUPPRESSOR OF MAX2 1-LIKE proteins form a regulatory module driving phloem development. Nat Commun 14: 2128. 10.1038/s41467-023-37790-5.

Wang F, Mei Z, Qi Y, Yan C, Hu Q, Wang J, Shi Y. 2011. Structure and mechanism of the hexameric MecA-ClpC molecular machine. Nature 471: 331–335. 10.1038/nature09780.

Wang L, Wang B, Jiang L, Liu X, Li X, Lu Z, Meng X, Wang Y, Smith SM, Li J. 2015. Strigolactone Signaling in Arabidopsis Regulates Shoot Development by Targeting D53-Like SMXL Repressor Proteins for Ubiquitination and Degradation. Plant Cell 27: 3128–3142. 10.1105/tpc.15.00605.

Wang L, Wang B, Yu H, Guo H, Lin T, Kou L, Wang A, Shao N, Ma H, Xiong G, et al. 2020a. Transcriptional regulation of strigolactone signalling in Arabidopsis. Nature 583: 277–281. 10.1038/s41586-020-2382-x.

Wang L, Xu Q, Yu H, Ma H, Li X, Yang J, Chu J, Xie Q, Wang Y, Smith SM, et al. 2020b. Strigolactone and Karrikin Signaling Pathways Elicit Ubiquitination and Proteolysis of SMXL2 to Regulate Hypocotyl Elongation in Arabidopsis. Plant Cell 32: 2251–2270. 10.1105/tpc.20.00140.

Wang M, Le Moigne M-A, Bertheloot J, Crespel L, Perez-Garcia M-D, Ogé L, Demotes-Mainard S, Hamama L, Davière J-M, Sakr S. 2019. BRANCHED1: A Key Hub of Shoot Branching. Front Plant Sci 10: 76. 10.3389/fpls.2019.00076.

Waters MT, Nelson DC. 2022. Karrikin perception and signalling. New Phytol. 10.1111/nph.18598.

White ARF, Mendez JA, Khosla A, Nelson DC. 2022. Rapid analysis of strigolactone receptor activity in a *Nicotiana benthamiana dwarf14* mutant. Plant Direct 6. 10.1002/pld3.389.

Wu Y-Y, Hou B-H, Lee W-C, Lu S-H, Yang C-J, Vaucheret H, Chen H-M. 2017. DCL2- and RDR6- dependent transitive silencing of SMXL4 and SMXL5 in Arabidopsis dcl4 mutants causes defective phloem transport and carbohydrate over-accumulation. Plant J 90: 1064–1078. 10.1111/tpj.13528.

Xu P, Hu J, Chen H, Cai W. 2023. SMAX1 interacts with DELLA protein to inhibit seed germination under weak light conditions via gibberellin biosynthesis in Arabidopsis. Cell Rep 42: 112740. 10.1016/j.celrep.2023.112740.

Yang T, Lian Y, Kang J, Bian Z, Xuan L, Gao Z, Wang X, Deng J, Wang C. 2020. The SUPPRESSOR of MAX2 1 (SMAX1)-Like SMXL6, SMXL7 and SMXL8 Act as Negative Regulators in Response to Drought Stress in Arabidopsis. Plant Cell Physiol 61: 1477–1492. 10.1093/pcp/pcaa066.

Yao R, Ming Z, Yan L, Li S, Wang F, Ma S, Yu C, Yang M, Chen L, Chen L, et al. 2016. DWARF14 is a non-canonical hormone receptor for strigolactone. Nature 536: 469–473. 10.1038/nature19073.

Yin L, Zander M, Huang S-SC, Xie M, Song L, Paola Saldierna Guzmán J, Hann E, Shanbhag BK, Ng S, Jain S, et al. 2023. Transcription Factor Dynamics in Cross-Regulation of Plant Hormone Signaling Pathways. bioRxiv 2023.03.07.531630. https://www.biorxiv.org/content/10.1101/2023.03.07.531630v1.abstract (Accessed April 18, 2024).

Zheng J, Hong K, Zeng L, Wang L, Kang S, Qu M, Dai J, Zou L, Zhu L, Tang Z, et al. 2020. Karrikin Signaling Acts Parallel to and Additively with Strigolactone Signaling to Regulate Rice Mesocotyl Elongation in Darkness. Plant Cell 32: 2780–2805. 10.1105/tpc.20.00123.

Zhou F, Lin Q, Zhu L, Ren Y, Zhou K, Shabek N, Wu F, Mao H, Dong W, Gan L, et al 2013. D14-SCF(D3)-dependent degradation of D53 regulates strigolactone signalling. Nature 504: 406–410. 10.1038/nature12878.

