## Supplemental Tables for "An N-terminal domain specifies developmental control by the SMAX1-LIKE family of transcriptional co-repressors in *Arabidopsis thaliana*"

**Supplemental Table 1.** Details of disorder prediction for SMAX1 and SMXL7 by D$^2$P$^2$

| | |
|---|---|
| **Protein** | ATSMAX1 |
| **Accession** | AT5G57710.1 |
| **Protein sequence*** | MRA**GLSTI**QQT**L**TPEAATVLNQSIA**EAARRN**HGQTTPLHVAATLLASPAGFLRRACIRSHPNSSHPLQCRA<br>LELCFSVALERLPT**ATTTPGNDPPISN**ALMAALKRAQAHQRRGCP**EQQQQ**PLLAVKVELEQLIISILDDPSV<br>SRVMREASFSSPAVKAT**IEQSLNNSVT**P**T**PIP**SVSSVGLNFRP**G**GGGPMTR**NSYLNPR**LQQNASSVQSG**<br>**VS**KNDDVERVMDILGRAKKKNPVLVGDSEPGRVIREILKKIEVGEVGNLAVKNSKVVSLEEISSDKALRIKEL<br>DGLLQTRLKNSDPIGGGGVILDLGDLKWLVEQPSSTQPPATVAVEIGRTAVVELRRLLEKFEGRLWFIGTA<br>TCETYLRCQVYHPSVETDWDLQAVSVAAKAPASGVFPRLANNLESFTPLKSFVPANRTLKCCPQCLQSYE<br>RELAEIDSVSSPEV**K**SEVAQPKQLPQWLLKAKPVDRLPQAKIEEVQKKWNDACVRLHPSFHNKNERIVPIP<br>VPITLTTSPYS**P**N**MLLRQPLQPKL**Q**PNRE**LRERVH**LKPMSPLVAEQAKKKSPPGSPVQTDLVLGR**AE**D**S<br>EKAGDVQVRDFLGCISSESVQNNNNISVLQKENLGNSLDIDLFKKLLKGMTEKVWWQNDAAAAVAATVSQ<br>CKLGNGKRRGVLSKGDVWLLFSGPDRVGKRKMVSALSSLVYGTNPIMIQLGSRQDAGDGNSSFRGKTAL<br>DKIAETVKRSPFSVILLEDIDEADMLVRGSIKQAMDRGRIRDSHGREISLGNVIFVMTASWHFAGTKTSFLD<br>NEAKLRDLASESWRLRLCMREKFGKRRASWLCSDEERLTKP**KKEH**GSG**LSFDLNQAA**DT**DDGSHNTSD**<br>**LTTDNDQD**EQGFSGKLSLQCVPFAFHDMVSRVDDAVAFRAVDFAAVRRRITETLSERFETIIGESLSVEVE<br>EEALQRILSGVWLGQTELEEWIEKAIVPVLSQLKARVSSSGTYGDCTVARLELDEDSGERNAGDLLPTTITL<br>AV |

* The **highlighted portions of sequence** are where there is 75% agreement between all predictors in the database for this region being disordered.

**Predicted Disordered Regions****

| Start | End |
|---|---|
| 4 | 8 |
| 12 | 12 |
| 26 | 31 |
| 86 | 98 |
| 117 | 121 |
| 161 | 170 |
| 172 | 172 |
| 176 | 186 |
| 188 | 194 |
| 202 | 214 |
| 441 | 441 |
| 509 | 509 |
| 511 | 521 |
| 523 | 526 |
| 533 | 562 |
| 565 | 565 |
| 819 | 822 |
| 826 | 834 |
| 837 | 853 |

* Regions over 75% agreement between all predictors in the database for this region being disordered.

**List of Disordered Regions by Predictor:**

| Predictor | Start | End |
|---|---|---|
| VLXT | 1 | 2 |
| VSL2b | 1 | 20 |
| PrDOS | 1 | 13 |
| PV2 | 1 | 39 |
| IUPred-S | 1 | 9 |
| Espritz-N | 1 | 13 |
| Espritz-X | 1 | 8 |
| IUPred-L | 3 | 7 |
| VLXT | 4 | 31 |
| IUPred-L | 12 | 12 |
| IUPred-L | 19 | 33 |
| VSL2b | 23 | 39 |
| PrDOS | 25 | 32 |
| Espritz-N | 26 | 37 |
| IUPred-S | 28 | 28 |

| | | |
|---|---|---|
| IUPred-S | 30 | 31 |
| IUPred-L | 35 | 39 |
| VLXT | 36 | 36 |
| PV2 | 48 | 48 |
| VSL2b | 56 | 68 |
| PV2 | 56 | 61 |
| PrDOS | 58 | 68 |
| Espritz-N | 58 | 66 |
| PV2 | 63 | 68 |
| PrDOS | 81 | 123 |
| Espritz-N | 81 | 98 |
| VSL2b | 82 | 129 |
| PV2 | 82 | 131 |
| VLXT | 84 | 104 |
| IUPred-L | 84 | 84 |
| IUPred-S | 86 | 105 |
| IUPred-L | 86 | 103 |
| Espritz-X | 88 | 90 |
| Espritz-N | 105 | 122 |
| IUPred-S | 107 | 116 |
| IUPred-L | 107 | 121 |
| VLXT | 117 | 126 |
| IUPred-S | 118 | 118 |
| VLXT | 133 | 139 |
| VSL2b | 134 | 135 |
| PV2 | 134 | 135 |
| VSL2b | 138 | 245 |
| PV2 | 138 | 237 |
| Espritz-N | 141 | 142 |
| IUPred-L | 148 | 149 |
| VLXT | 149 | 254 |
| Espritz-N | 150 | 194 |
| IUPred-L | 157 | 157 |
| PrDOS | 160 | 219 |
| IUPred-L | 160 | 170 |
| IUPred-S | 170 | 170 |
| IUPred-S | 172 | 172 |
| IUPred-L | 172 | 172 |
| IUPred-L | 175 | 186 |
| IUPred-S | 180 | 216 |
| IUPred-L | 188 | 214 |
| Espritz-N | 202 | 218 |
| IUPred-L | 223 | 224 |
| IUPred-S | 225 | 225 |
| Espritz-N | 229 | 242 |
| IUPred-L | 233 | 233 |
| IUPred-L | 237 | 237 |
| PV2 | 240 | 241 |
| PV2 | 244 | 244 |
| VSL2b | 254 | 254 |
| PrDOS | 255 | 266 |
| VSL2b | 256 | 260 |
| VSL2b | 262 | 280 |
| PrDOS | 272 | 283 |
| VLXT | 273 | 292 |
| PV2 | 277 | 280 |
| Espritz-N | 294 | 303 |

| | | |
|---|---|---|
| VSL2b | 295 | 297 |
| PV2 | 301 | 304 |
| PV2 | 315 | 329 |
| VSL2b | 318 | 325 |
| Espritz-N | 319 | 321 |
| PrDOS | 320 | 334 |
| VLXT | 321 | 339 |
| IUPred-S | 326 | 326 |
| PrDOS | 383 | 414 |
| PV2 | 387 | 389 |
| VSL2b | 413 | 413 |
| VSL2b | 415 | 415 |
| VSL2b | 417 | 471 |
| PV2 | 418 | 472 |
| VLXT | 424 | 442 |
| PrDOS | 431 | 470 |
| Espritz-N | 435 | 450 |
| IUPred-L | 440 | 441 |
| IUPred-S | 443 | 443 |
| VLXT | 452 | 471 |
| IUPred-L | 466 | 467 |
| PV2 | 474 | 476 |
| PrDOS | 482 | 606 |
| Espritz-N | 484 | 526 |
| PV2 | 490 | 490 |
| VLXT | 497 | 567 |
| IUPred-L | 498 | 500 |
| PV2 | 499 | 575 |
| VSL2b | 502 | 575 |
| IUPred-L | 505 | 505 |
| IUPred-L | 507 | 509 |
| IUPred-S | 511 | 511 |
| IUPred-L | 511 | 521 |
| IUPred-S | 513 | 563 |
| IUPred-L | 523 | 562 |
| Espritz-N | 533 | 557 |
| Espritz-X | 541 | 561 |
| Espritz-N | 560 | 570 |
| IUPred-S | 565 | 565 |
| IUPred-L | 565 | 565 |
| VLXT | 573 | 573 |
| PV2 | 582 | 592 |
| VSL2b | 584 | 592 |
| Espritz-N | 586 | 593 |
| IUPred-L | 592 | 592 |
| IUPred-L | 594 | 594 |
| VSL2b | 596 | 598 |
| PV2 | 596 | 598 |
| Espritz-N | 597 | 602 |
| PrDOS | 637 | 652 |
| VLXT | 638 | 649 |
| VSL2b | 642 | 643 |
| VLXT | 656 | 656 |
| VSL2b | 665 | 666 |
| VLXT | 669 | 688 |
| PV2 | 680 | 681 |
| PV2 | 683 | 687 |

| | | |
|---|---|---|
| PrDOS | 687 | 705 |
| IUPred-S | 688 | 689 |
| Espritz-N | 688 | 701 |
| VSL2b | 690 | 702 |
| PV2 | 690 | 702 |
| Espritz-X | 693 | 695 |
| IUPred-S | 695 | 695 |
| IUPred-S | 697 | 698 |
| VLXT | 700 | 706 |
| PV2 | 707 | 709 |
| VLXT | 715 | 722 |
| VLXT | 729 | 747 |
| PV2 | 730 | 730 |
| VSL2b | 739 | 750 |
| IUPred-L | 743 | 743 |
| PV2 | 744 | 746 |
| Espritz-N | 746 | 750 |
| PrDOS | 767 | 872 |
| VSL2b | 779 | 779 |
| PV2 | 784 | 784 |
| Espritz-N | 784 | 788 |
| VLXT | 785 | 820 |
| VSL2b | 792 | 859 |
| PV2 | 792 | 794 |
| PV2 | 797 | 860 |
| Espritz-N | 798 | 865 |
| Espritz-X | 801 | 864 |
| IUPred-S | 820 | 855 |
| IUPred-L | 820 | 822 |
| IUPred-L | 826 | 834 |
| IUPred-L | 837 | 853 |
| VLXT | 840 | 854 |
| VLXT | 901 | 927 |
| VSL2b | 913 | 917 |
| PV2 | 916 | 916 |
| VSL2b | 952 | 956 |
| PV2 | 953 | 953 |
| PrDOS | 954 | 962 |
| Espritz-N | 954 | 959 |
| VLXT | 962 | 981 |
| PV2 | 966 | 982 |
| PrDOS | 968 | 990 |
| VSL2b | 969 | 979 |
| Espritz-N | 973 | 978 |
| Espritz-X | 975 | 981 |
| IUPred-S | 984 | 990 |
| Espritz-X | 987 | 990 |
| VSL2b | 988 | 990 |
| PV2 | 988 | 990 |

| | |
|---|---|
| **Protein** | ATSMXL7 |
| **Accession** | AT2G29970.1 |
| **Protein sequence*** | **MPTPVTT**ARQCLTEETARALDDAVSVARRRSHAQTTSLHAVSGLLTMPSSILREVCISRAAHNTPYSSRLQ<br>FRALELCVGVSLDRLP**SSKSTPTTTVEEDPPVS**NSLMAA**IKRSQ**ATQRRHPETYHLHQIHGNNNTETTSVL<br>KVELKYFILSILDDPIVSRVFGEAGFRSTDIKLDVLHPPVTSQFSSRFTSRSRIPPLFLCNLPESDSGRVRFG<br>FPFGDLDENCRRIGEVLARKDKKNPLLVGVCGVEALKTFTDSINRGKFGFLPLEISGLSVVSIKISEVLVDGS<br>RIDIKFDDLGRLKSGMVLNLGELKVLASDVFSVDVIEKFVLKLADLLKLHREKLWFIGSVSSNETYLKLIERF<br>PTIDKDWNLHLLPITSSSQGLYPKSSLMGSFVPFGGFFSSTSDFRIPSSSSMNQTLPRCHLCNEKYEQEVT<br>AFAKSGSMIDDQCSEKLPSWLRNVEHEHEKG**NLGKVKDD**PNVLASRIPALQKKWDDICQRIHQTPAFPKL<br>SFQPVRPQFPLQLG**SSSQTK**MSL**GS**PTEKIVCTRTSESFQG**MVALPQNPPHQPGLSV**KISK**PKHTEDLS**S<br>STTNSPLSFVTTDLGLGTIYASKN**QEPS**TP**VSVE**RRDFEVIKEKQLLSASRYCKDFKSLRELLSRKVGFQN<br>EAVNAISEIVCGYRDESRRRNNHVATTSNVWLALLGPDKAGKKKVALALAEVFCGGQDNFICVDFKSQDS<br>LDDRFRGKTVVDYIAGEVARRADSVVFIENVEKAEFPDQIRLSEAMRTGKLRDSHGREISMKNVIVVATISG<br>SDKASDCHVLEEPVKYSEERVLNAKNWTLQIKLADTS**NVNKNGPNKRRQEEAETE**VTELRALKSQRSFLD<br>LNLPVDEIEAN**EDE**AYTMSENTEAWLEDFVEQVDGKVTFKLIDFDELAKNIKRNILSLFHLSFGPETHLEIEN<br>DVILKILAALRWSSDEEKTFDQWLQTVLAPSFAKARQKCVPAAPFSVKLVASRESPAEE**ETT**GIQQFPARV<br>EVI |

* The **highlighted portions of sequence** are where there is 75% agreement between all predictors in the database for this region being disordered.

**Predicted Disordered Regions****

| Start | End |
|---|---|
| 1 | 7 |
| 88 | 104 |
| 111 | 115 |
| 464 | 471 |
| 517 | 522 |
| 526 | 527 |
| 544 | 559 |
| 564 | 571 |
| 597 | 600 |
| 603 | 606 |
| 823 | 840 |
| 867 | 869 |
| 988 | 990 |

** Regions over 75% agreement between all predictors in the database for this region being disordered.

**List of Disordered Regions by Predictor:**

| Predictor | Start | End |
|---|---|---|
| VLXT | 1 | 9 |
| VSL2b | 1 | 43 |
| PrDOS | 1 | 11 |
| PV2 | 1 | 43 |
| IUPred-S | 1 | 10 |
| IUPred-L | 1 | 14 |
| Espritz-N | 1 | 6 |
| Espritz-X | 1 | 7 |
| IUPred-S | 12 | 12 |
| IUPred-L | 22 | 33 |
| Espritz-N | 32 | 34 |
| IUPred-L | 35 | 40 |
| VLXT | 36 | 45 |
| PrDOS | 54 | 69 |
| Espritz-N | 61 | 66 |
| PV2 | 64 | 68 |
| PV2 | 71 | 72 |
| PV2 | 75 | 78 |
| PV2 | 80 | 138 |
| Espritz-N | 82 | 104 |
| VSL2b | 83 | 126 |
| VLXT | 86 | 117 |
| PrDOS | 86 | 143 |

| | | |
|---|---|---|
| Espritz-X | 87 | 97 |
| IUPred-L | 88 | 115 |
| IUPred-S | 89 | 131 |
| Espritz-N | 111 | 137 |
| IUPred-L | 118 | 132 |
| VSL2b | 131 | 138 |
| PrDOS | 164 | 222 |
| PV2 | 174 | 177 |
| VSL2b | 181 | 215 |
| PV2 | 181 | 207 |
| Espritz-N | 191 | 195 |
| Espritz-N | 207 | 209 |
| PV2 | 209 | 211 |
| VSL2b | 217 | 217 |
| PV2 | 222 | 222 |
| VLXT | 227 | 237 |
| PrDOS | 284 | 292 |
| PV2 | 317 | 317 |
| PV2 | 324 | 327 |
| Espritz-N | 377 | 388 |
| VLXT | 378 | 383 |
| VSL2b | 378 | 419 |
| PrDOS | 378 | 416 |
| PV2 | 380 | 381 |
| PV2 | 383 | 384 |
| PV2 | 389 | 391 |
| PV2 | 393 | 399 |
| Espritz-N | 396 | 417 |
| PV2 | 401 | 406 |
| PV2 | 409 | 417 |
| PV2 | 419 | 419 |
| VSL2b | 427 | 432 |
| PV2 | 427 | 432 |
| PrDOS | 431 | 482 |
| Espritz-N | 436 | 444 |
| PV2 | 438 | 438 |
| VSL2b | 440 | 445 |
| PV2 | 440 | 477 |
| VLXT | 450 | 453 |
| IUPred-L | 452 | 453 |
| VSL2b | 455 | 471 |
| IUPred-L | 456 | 456 |
| Espritz-N | 458 | 473 |
| IUPred-L | 460 | 474 |
| IUPred-S | 463 | 465 |
| VLXT | 464 | 479 |
| IUPred-S | 467 | 469 |
| PV2 | 480 | 480 |
| PV2 | 483 | 484 |
| Espritz-N | 485 | 485 |
| PrDOS | 494 | 585 |
| Espritz-N | 494 | 532 |
| VSL2b | 496 | 608 |
| PV2 | 496 | 606 |
| IUPred-S | 501 | 501 |
| IUPred-L | 501 | 501 |
| IUPred-L | 510 | 511 |

| | | |
|---|---|---|
| IUPred-S | 513 | 516 |
| IUPred-L | 513 | 513 |
| IUPred-L | 515 | 522 |
| VLXT | 517 | 527 |
| IUPred-S | 518 | 524 |
| IUPred-L | 524 | 529 |
| IUPred-S | 526 | 531 |
| IUPred-S | 534 | 537 |
| Espritz-N | 535 | 581 |
| IUPred-S | 543 | 576 |
| IUPred-L | 543 | 576 |
| VLXT | 544 | 559 |
| Espritz-X | 550 | 553 |
| Espritz-X | 564 | 571 |
| Espritz-N | 585 | 587 |
| PrDOS | 591 | 628 |
| VLXT | 592 | 616 |
| IUPred-S | 593 | 593 |
| IUPred-L | 593 | 593 |
| Espritz-N | 594 | 607 |
| IUPred-L | 597 | 600 |
| IUPred-S | 599 | 600 |
| IUPred-L | 603 | 607 |
| PV2 | 613 | 613 |
| VSL2b | 614 | 615 |
| PV2 | 615 | 615 |
| VLXT | 626 | 633 |
| PrDOS | 655 | 672 |
| VSL2b | 660 | 666 |
| Espritz-N | 660 | 663 |
| PV2 | 662 | 662 |
| PV2 | 664 | 667 |
| VLXT | 678 | 681 |
| VSL2b | 682 | 686 |
| PV2 | 685 | 688 |
| PrDOS | 710 | 725 |
| VSL2b | 713 | 716 |
| PV2 | 716 | 717 |
| PrDOS | 731 | 733 |
| VLXT | 736 | 772 |
| PV2 | 749 | 753 |
| Espritz-N | 757 | 771 |
| VSL2b | 760 | 771 |
| IUPred-L | 760 | 760 |
| PV2 | 762 | 768 |
| IUPred-L | 765 | 766 |
| IUPred-S | 766 | 767 |
| PV2 | 770 | 770 |
| Espritz-N | 784 | 789 |
| PrDOS | 785 | 884 |
| VSL2b | 786 | 790 |
| PV2 | 786 | 787 |
| PV2 | 789 | 791 |
| IUPred-L | 794 | 794 |
| PV2 | 814 | 873 |
| VSL2b | 818 | 874 |
| Espritz-N | 818 | 837 |

| | | |
|---|---|---|
| VLXT | 820 | 850 |
| IUPred-S | 823 | 843 |
| IUPred-L | 823 | 841 |
| Espritz-X | 826 | 840 |
| VLXT | 854 | 876 |
| Espritz-N | 862 | 876 |
| IUPred-L | 865 | 865 |
| IUPred-S | 867 | 871 |
| IUPred-L | 867 | 869 |
| IUPred-L | 872 | 872 |
| IUPred-S | 874 | 874 |
| VLXT | 925 | 925 |
| VSL2b | 944 | 945 |
| PV2 | 961 | 1002 |
| VSL2b | 966 | 1002 |
| VLXT | 977 | 994 |
| PrDOS | 981 | 1002 |
| Espritz-N | 982 | 990 |
| Espritz-X | 983 | 1002 |
| IUPred-S | 988 | 991 |
| IUPred-L | 989 | 990 |
| IUPred-S | 996 | 1002 |
| VLXT | 1001 | 1001 |

### Supplemental Table 2. TFs used in Y2H assay

| Arabidopsis TF library number[1] | TF family | Accession | Gene name | Interaction with SMAX1/SMXL7 | References |
|---|---|---|---|---|---|
| DEST-U18-D06 | ABI3-VP1 | AT3G24650.1 | ABI3 | No | Seed dormancy[2,3] |
| DEST-U13-F01 | ALFIN-like | AT2G02470.2 | AL6 | No | Root hair development[4] |
| DEST-U17-C04 | AP2-EREBP | AT5G10510.1 | AIL6 | No | *rac*-GR24 responsive Differently Accessible Region (DAR)[5] |
| DEST-U15-G10 | AP2-EREBP | AT1G24590.1 | BOL | SMAX1 | Germination and embryo morphogenesis [2] |
| DEST-U20-C11 | AP2-EREBP | AT1G12980.1 | DRN | SMAX1 | Germination and embryo morphogenesis [2] |
| DEST-U04-A08 | AP2-EREBP | AT3G23240.1 | ERF1 | No | Photomorphogenesis[6] |
| DEST-U05-A09 | AP2-EREBP | AT4G34410.1 | ERF109 | No | $GR24^{4DO}$-responsive gene[7] |
| DEST-U07-F04 | AP2-EREBP | AT1G28370.1 | ERF11 | No | *rac*-GR24 responsive gene[5] |
| DEST-U16-H12 | AP2-EREBP | AT5G25810.1 | ERF40 | No | *rac*-GR24 responsive DAR[5] |
| DEST-U05-E05 | AP2-EREBP | AT5G65130.1 | ERF57 | No | *rac*-GR24 responsive gene[5] |
| DEST-U17-D07 | AP2-EREBP | AT4G17490.1 | ERF6 | No | *rac*-GR24 responsive DAR[5] |
| DEST-U01-H06 | AP2-EREBP | AT1G64380.1 | ERF61 | SMAX1 and SMXL7 | SMXL6-targeted and *rac*-GR24-responsive gene[5,7] |
| DEST-U05-C06 | AP2-EREBP | AT3G16770.1 | ERF72 | No | SMXL6-targeted[7] |
| DEST-U05-F07 | AP2-EREBP | AT5G44210.1 | ERF9 | No | SMXL6-targeted[7] |
| DEST-U03-E05 | AP2-EREBP | AT5G25190.1 | ESE3 | No | SMXL6-targeted[7] |
| DEST-U18-D08 | AP2-EREBP | AT3G20840.1 | PLT1 | No | Germination and embryo morphogenesis [2] |
| DEST-U05-E01 | AP2-EREBP | AT1G51190.1 | PLT2 | No | Germination and embryo morphogenesis [2] |
| DEST-U19-G05 | ARF | AT1G59750.1 | ARF1 | No | *rac*-GR24 responsive gene[5] |
| DEST-U07-B08 | ARF | AT3G61830.1 | ARF18 | No | *rac*-GR24 responsive gene[5] |
| DEST-U05-H03 | AUX/IAA | AT3G15540.1 | IAA19 | No | SMXL6-targeted[7] |
| DEST-U18-C01 | AUX/IAA | AT4G32280.1 | IAA29 | No | Cell elongation regulation[8] |
| DEST-U04-F12 | AUX/IAA | AT5G43700.1 | IAA4 | No | *rac*-GR24 responsive gene[5] |
| DEST-U02-D06 | AUX/IAA | AT1G15580.1 | IAA5 | No | SMXL6-targeted[7] |
| DEST-U04-F08 | BES1 | AT3G50750.1 | BEH1 | No | Arabidopsis homolog of OsBES1[9], Photomorphogenesis and seed germination[10] |
| DEST-U17-F04 | BES1 | AT4G36780.2 | BEH2 | No | Arabidopsis homolog of OsBES1[9], Photomorphogenesis and seed germination[10] |
| DEST-U06-F06 | BES1 | AT4G18890.1 | BEH3 | No | Arabidopsis homolog of OsBES1[9] |
| DEST-U16-H10 | BES1 | AT1G19350.1 | BES1 | No | Arabidopsis homolog of OsBES1[9], Photomorphogenesis and seed germination[10] |
| DEST-U10-C09 | BES1 | AT1G75080.1 | BZR1 | No | Photomorphogenesis and seed germination[10] |
| DEST-U15-G04 | bHLH | AT1G68810.1 | ABS5/T5L1 | SMAX1 | Germination[2] |
| DEST-U12-E12 | bHLH | AT2G28160.1 | FIT | No | GR244DO-responsive and targeted by SMXL6[7] |
| DEST-U01-D07 | bHLH | AT5G67060.1 | HEC1 | No | GR244DO-responsive and targeted by SMXL6[7] |

| | | | | | |
|---|---|---|---|---|---|
| DEST-U02-F12 | bHLH | AT4G30180.1 | HLH4 | No | Cell elongation and anthocyanin accumulation[11] |
| DEST-U19-A07 | bHLH | AT2G27230.1 | LHW | No | Germination and Seedling development[2] |
| DEST-U19-B01 | bHLH | AT1G32640.1 | MYC2 | No | Photomorphogenesis[12] |
| DEST-U13-A07 | bHLH | AT5G46760.1 | MYC3 | No | Seed germination regulation[13], *rac*-GR24-responsive gene[5] |
| DEST-U13-A05 | bHLH | AT4G17880.1 | MYC4 | No | Seed germination regulation[13] |
| DEST-U01-C06 | bHLH | AT2G20180.2 | PIF1 | No | Photomorphogenesis[14] |
| DEST-U15-F08 | bHLH | AT2G43010.1 | PIF4 | No | Photomorphogenesis[10,15], thermomorphogenesis[16] |
| DEST-U20-E03 | bHLH | AT3G59060.2 | PIF5 | No | Photomorphogenesis[14] |
| DEST-U09-H06 | bHLH | AT5G61270.1 | PIF7 | No | Photomorphogenesis and thermomorphogenesis[17] |
| DEST-U07-A04 | bHLH | AT4G00050.1 | PIF8 | No | Photomorphogenesis[18], potential genetic interaction downstream of PhyB[16] |
| DEST-U04-G09 | bHLH | AT1G66470.1 | RHD6 | No | Root hair development[4] |
| DEST-U09-H05 | bHLH | AT5G37800.1 | RSL1 | No | Root hair development[4] |
| DEST-U01-C01 | bHLH | AT1G09530.1 | PIF3 | SMAX1 | Photomorphogenesis[10,15], thermomorphogenesis[19] |
| DEST-U06-H12 | bHLH | AT4G36930.1 | SPT | SMAX1 | Seed dormancy[2] |
| DEST-U03-F05 | bHLH | AT3G25710.1 | TMO5 | SMAX1 | Seed development[2] |
| DEST-U06-B10 | bZIP | AT1G49720.1 | ABF1 | No | Germination[3] |
| DEST-U06-B05 | bZIP | AT1G45249.1 | ABF2 | No | Seed dormancy[2,3] |
| DEST-U18-E06 | bZIP | AT4G34000.1 | ABF3 | No | Germination[3] |
| DEST-U02-H03 | bZIP | AT3G19290.1 | ABF4 | No | Germination[3] |
| DEST-U06-B06 | bZIP | AT2G36270.1 | ABI5 | No | Seed dormancy[2,3] |
| DEST-U09-F02 | bZIP | AT4G01120.1 | bZIP54/GBF2 | No | Seed development[2] |
| DEST-U03-D06 | bZIP | AT3G44460.1 | bZIP67 | No | Seed dormancy[2,3] |
| DEST-U16-B08 | bZIP | AT5G11260.1 | HY5 | No | Photomorphogenesis[15], seed germination[10] |
| DEST-U02-B02 | bZIP | AT3G17609.4 | HYH | No | Photomorphogenesis[15] |
| DEST-U12-F01 | bZIP | AT1G22070.1 | TGA3/bZIP22 | SMAX1 | Seed dormancy regulation[3] |
| DEST-U01-F06 | C2C2-CO-like | AT2G47890.1 | BBX11 | No | Photomorphogenesis, thermomorphogenesis, and flowering [10,20] |
| DEST-U08-A07 | C2C2-CO-like | AT2G24790.1 | BBX4 | No | Photomorphogenesis, flowering[10,20] |
| DEST-U11-D09 | C2C2-DOF | AT3G61850.1 | DAG1 | No | Germination regulation[3] |
| DEST-U05-F02 | C2C2-YABBY | AT2G45190.1 | YAB1/FIL | No | Germination and embryo morphogenesis [2] |
| DEST-U01-E05 | C2C2-YABBY | AT1G08465.1 | YAB2 | No | Germination and embryo morphogenesis [2] |
| DEST-U04-B10 | C2C2-YABBY | AT4G00180.1 | YAB3 | No | Germination and embryo morphogenesis [2] |
| DEST-U07-G05 | C2H2 | AT2G34500.1 | FZF | No | *rac*-GR24 responsive DAR[5] |
| DEST-U06-G09 | C2H2 | AT1G10480.1 | ZFP5 | No | Root hair development[4] |
| DEST-U17-B02 | EIL | AT2G27050.1 | EIL1 | No | Photomorphogenesis[6], Root hair development[4] |
| DEST-U19-B12 | EIL | AT3G20770.1 | EIN3 | No | Photomorphogenesis[6], Root hair development[4] |

| | | | | | |
|---|---|---|---|---|---|
| DEST-U06-C01 | G2-like | AT5G16560.1 | KAN1 | No | Germination and embryo morphogenesis[2], Leaf development[21] |
| DEST-U16-B05 | G2-like | AT1G32240.1 | KAN2 | No | Germination and embryo morphogenesis[2], leaf development[22] |
| DEST-U06-H02 | G2-like | AT5G42630.1 | KAN4 | No | Germination and embryo morphogenesis[2] |
| DEST-U17-B04 | GNAT | AT4G37580.1 | COP3/HLS1 | No | Thermomorphogenesis[23] |
| DEST-U04-H01 | GRAS | AT2G01570.1 | RGA | SMAX1 | Photomorphogenesis and seed germination[10,24], seed dormancy[2] |
| DEST-U12-E08 | GRAS | AT1G66350.1 | RGL1 | SMAX1 and SMXL7 | Seed dormancy and germination, photomorphogenesis[10] |
| DEST-U04-C04 | GRAS | AT1G14920.1 | GAI | SMAX1 and SMXL7 | Photomorphogenesis and seed germination[10,24] |
| DEST-U07-F03 | GRAS | AT5G17490.1 | RGL3 | SMAX1 and SMXL7 | Photomorphogenesis and seed germination[10,24], seed dormancy[2] |
| DEST-U16-C02 | GRF | AT2G22840.1 | GRF1 | No | Arabidopsis homolog of OsGRF4[25] |
| DEST-U12-F05 | GRF | AT4G37740.1 | GRF2 | No | Arabidopsis homolog of OsGRF4[25] |
| DEST-U11-A10 | GRF | AT2G36400.1 | GRF3 | No | Arabidopsis homolog of OsGRF4[25] |
| DEST-U17-B10 | GRF | AT3G52910.1 | GRF4 | No | Arabidopsis homolog of OsGRF4[25] |
| DEST-U16-H03 | GRF | AT5G53660.1 | GRF7 | No | Arabidopsis homolog of OsGRF4[25] |
| DEST-U16-C04 | GRF | AT2G45480.1 | GRF9 | No | Arabidopsis homolog of OsGRF4[25] |
| DEST-U04-C06 | HB | AT3G01470.1 | ATHB1 | No | Hypocotyl elongation[26], *rac*-GR24-responsive gene[5] |
| DEST-U18-D04 | HB | AT2G34710.1 | ATHB14/PHB | SMAX1 | Germination and embryo morphogenesis[2] |
| DEST-U19-G03 | HB | AT1G52150.1 | ATHB15/CNA | SMAX1 and SMXL7 | Germination and embryo morphogenesis[2] |
| DEST-U19-G04 | HB | AT1G30490.1 | ATHB9/PHV | SMAX1 and SMXL7 | Germination and embryo morphogenesis[2] |
| DEST-U19-G09 | HB | AT1G79840.1 | GL2 | No | Root hair development[4] |
| DEST-U04-G11 | HB | AT3G60390.1 | HAT3 | No | Germination and embryo morphogenesis[2] |
| DEST-U03-E09 | HB | AT4G16780.1 | HAT4/ATHB2 | No | Germination and embryo morphogenesis[2] |
| DEST-U14-A07 | HB | AT4G04890.1 | PDF2 | SMAX1 and SMXL7 | Germination and embryo morphogenesis[2] |
| DEST-U01-D09 | HB | AT5G59340.1 | WOX2 | No | Germination and embryo morphogenesis[2] |
| DEST-U01-D10 | HB | AT5G45980.1 | WOX8 | SMAX1 and SMXL7 | Germination and embryo morphogenesis[2] |
| DEST-U01-D10 | HB | AT2G33880.1 | WOX9 | SMAX1 and SMXL7 | Germination and embryo morphogenesis[2] |
| DEST-U01-H02 | HB | AT2G44910.1 | ATHB4 | SMAX1 | Germination and embryo morphogenesis[2] |
| DEST-U14-D08 | HSF | AT3G22830.1 | HSFA6B | No | GR24$^{4DO}$-responsive gene[7] |
| DEST-U16-B06 | HSF | AT1G46264.1 | HSFB4 | No | *rac*-GR24 responsive DAR[5] |
| DEST-U19-E12 | JUMONJI | AT1G09060.1 | JMJ24 | No | Arabidopsis homolog of OsGRF4[25] |
| DEST-U09-G11 | JUMONJI | AT3G20810.1 | JMJ30 | No | Post-germination regulation[3] |
| DEST-U02-F05 | LOB/AS2 | AT2G42430.1 | LBD16 | No | GR24$^{4DO}$-responsive gene[7] |
| DEST-U03-F04 | MYB | AT5G45420.1 | maMYB | No | Root hair development[4] |
| DEST-U06-B01 | MYB | AT3G23250.1 | MYB15 | No | *rac*-GR24 responsive gene[5] |
| DEST-U17-F01 | MYB | AT1G56650.1 | PAP1 | No | GR24$^{4DO}$-responsive gene[7] |
| DEST-U12-H05 | MYB-related | AT5G17300.1 | RVE1 | No | Seed dormancy and germination[3,27] |

| | | | | | |
|---|---|---|---|---|---|
| DEST-U05-H09 | MYB-related | AT1G01520.1 | RVE3 | No | Seed dormancy, germination and embryo morphogenesis[2] |
| DEST-U06-F12 | NAC | AT3G15170.1 | CUC1 | No | Germination and embryo morphogenesis[2], leaf development [21] |
| DEST-U06-C02 | NAC | AT5G53950.1 | CUC2 | No | Germination and embryo morphogenesis[2], leaf development [21] |
| DEST-U13-B03 | NAC | AT1G76420.1 | CUC3 | No | Germination and embryo morphogenesis[2] |
| DEST-U03-G02 | NAC | AT3G49530.1 | NAC062 | No | *rac*-GR24 responsive gene[5] |
| DEST-U06-G05 | NAC | AT5G62380.1 | NAC101 | No | SMXL6-targeted[7] |
| DEST-U14-C09 | Orphans | AT2G21320.1 | BBX18 | No | Photomorphogenesis[10] |
| DEST-U05-A07 | Orphans | AT4G38960.1 | BBX19 | No | Photomorphogenesis and flowering[10,20] |
| DEST-U01-G04 | Orphans | AT4G39070.1 | BBX20 | No | Photomorphogenesis[10,20] |
| DEST-U18-H01 | Orphans | AT1G75540.1 | BBX21 | No | Seed germination[10,20] |
| DEST-U10-D03 | Orphans | AT1G78600.1 | BBX22 | No | Photomorphogenesis[10,20] |
| DEST-U03-B06 | Orphans | AT4G10240.1 | BBX23 | No | Photomorphogenesis and thermomorphogenesis[10,20] |
| DEST-U02-H01 | Orphans | AT1G06040.1 | BBX24 | No | Photomorphogenesis[10,20] |
| DEST-U02-H09 | Orphans | AT2G31380.1 | BBX25 | No | Photomorphogenesis[10,20] |
| DEST-U12-D10 | Orphans | AT4G27310.1 | BBX28 | No | Photomorphogenesis[10,20] |
| DEST-U17-E01 | Orphans | AT5G54470.1 | BBX29 | No | Photomorphogenesis[10,20] |
| DEST-U08-E06 | Orphans | AT4G15248.1 | BBX30 | SMAX1 | Photomorphogenesis[10,20] |
| DEST-U14-B09 | Orphans | AT3G21890.1 | BBX31 | No | Photomorphogenesis[10,20] |
| DEST-U20-G12 | Orphans | AT3G23150.1 | ETR2 | No | $GR24^{4DO}$-responsive gene[7] |
| DEST-U19-F06 | Orphans | AT4G21430.1 | JMJ28 | No | Arabidopsis homolog of OsGRF4[25] |
| DEST-U14-G05 | Orphans | AT1G74890.1 | ARR15 | No | Seed development[14] |
| DEST-U14-G06 | Orphans | AT2G41310.1 | ARR8 | No | SMXL6-targeted[7] |
| DEST-U19-A05 | SBP | AT2G47070.1 | SPL1 | No | *Arabidopsis* homolog of OsIPA1[28] |
| DEST-U03-H07 | SBP | AT1G27370.1 | SPL10 | No | *Arabidopsis* homolog of OsIPA1[28] |
| DEST-U03-C08 | SBP | AT1G27360.1 | SPL11 | No | *Arabidopsis* homolog of OsIPA1[28] |
| DEST-U19-E05 | SBP | AT3G60030.1 | SPL12 | No | *Arabidopsis* homolog of OsIPA1[28] |
| DEST-U11-G04 | SBP | AT5G50570.1 | SPL13 | No | *Arabidopsis* homolog of OsIPA1[28] |
| DEST-U21-A02 | SBP | AT1G20980.1 | SPL14 | No | *Arabidopsis* homolog of OsIPA1[28] |
| DEST-U02-E03 | SBP | AT3G57920.1 | SPL15 | No | *Arabidopsis* homolog of OsIPA1[28] |
| DEST-U06-H04 | SBP | AT5G43270.1 | SPL2 | No | *Arabidopsis* homolog of OsIPA1[28] |
| DEST-U03-C12 | SBP | AT1G53160.1 | SPL4 | No | *Arabidopsis* homolog of OsIPA1[28] |
| DEST-U04-E09 | SBP | AT3G15270.1 | SPL5 | No | *Arabidopsis* homolog of OsIPA1[28] |
| DEST-U04-E08 | SBP | AT1G69170.1 | SPL6 | No | *Arabidopsis* homolog of OsIPA1[28] |
| DEST-U19-G10 | SBP | AT5G18830.1 | SPL7 | No | *Arabidopsis* homolog of OsIPA1[28] |
| DEST-U11-A06 | SBP | AT1G02065.1 | SPL8 | No | *Arabidopsis* homolog of OsIPA1[28] |

| | | | | | |
|---|---|---|---|---|---|
| DEST-U04-H02 | SBP | AT2G42200.1 | SPL9 | No | *Arabidopsis* homolog of OsIPA1[28] |
| DEST-U17-B01 | SIGMA70-like | AT3G53920.1 | SIG3 | No | Anterograde signals downstream of PIFs to regulate photoresponses[29] |
| DEST-U05-C04 | TCP | AT2G37000.1 | TCP (AT2G37000) | SMAX1 | member of TCP family |
| DEST-U01-E01 | TCP | AT5G41030.1 | TCP (AT5G41030) | No | member of TCP family |
| DEST-U07-G07 | TCP | AT2G31070.1 | TCP10 | SMAX1 and SMXL7 | Hypocotyl and leaf[30,31], member of TCP family |
| DEST-U17-E11 | TCP | AT1G68800.2 | TCP12/BRC2 | No | Branching[32,33], member of TCP family |
| DEST-U09-E02 | TCP | AT3G02150.2 | TCP13 | SMAX1 and SMXL7 | Hypocotyl and leaf[34,35], member of TCP family |
| DEST-U04-A09 | TCP | AT3G47620.1 | TCP14 | SMAX1 and SMXL7 | Hypocotyl and leaf petiole[36,37], member of TCP family |
| DEST-U15-D12 | TCP | AT3G45150.1 | TCP16 | SMAX1 and SMXL7 | Leaf development[38], member of TCP family |
| DEST-U01-E03 | TCP | AT5G08070.1 | TCP17 | SMAX1 | Hypocotyl elongation[35], member of TCP family |
| DEST-U15-E01 | TCP | AT3G18550.1 | TCP18/BRC1 | SMAX1 and SMXL7 | Branching[39–41], member of TCP family |
| DEST-U18-C02 | TCP | AT5G51910.1 | TCP19 | SMAX1 and SMXL7 | Member of TCP family, leaf senescence[42] |
| DEST-U20-D12 | TCP | AT5G08330.1 | TCP21 | SMAX1 | Hypocotyl elongation and leaf petiole[37], member of TCP family |
| DEST-U14-D09 | TCP | AT1G31210.1 | TCP24 | No | Hypocotyl elongation and leaf development[30,31], member of TCP family |
| DEST-U06-B07 | TCP | AT5G60970.1 | TCP5 | SMAX1 | Hypocoty elongation[35,43], member of TCP family |
| DEST-U20-B12 | TCP | AT5G23280.1 | TCP7 | SMAX1 and SMXL7 | Hypocotyl elongation and leaf petiole[37], member of TCP family |
| DEST-U07-C02 | TCP | AT1G58100.1 | TCP8 | SMAX1 and SMXL7 | Hypocotyl elongation and leaf petiole[37], member of TCP family |
| DEST-U07-F11 | TCP | AT2G45680.1 | TCP9 | SMAX1 and SMXL7 | Member of TCP family, leaf development[42] |
| DEST-U19-B11 | WRKY | AT5G56270.1 | WRKY2 | No | Germination and embryo morphogenesis[2] |
| DEST-U14-A02 | WRKY | AT2G38470.1 | WRKY33 | No | Phosphate-deficiency induced root architecture regulation[44] |
| DEST-U09-C09 | WRKY | AT4G04450.1 | WRKY42 | No | *rac*-GR24 responsive DAR[5] |
| DEST-U05-D04 | WRKY | AT5G13080.1 | WRKY75 | No | Root hair development[4] |

### Supplemental Table 3. Primers

I. Primers for cloning of pGWBcitr-SMAX1pro and pGWBcitr-SMXL7pro
(Backbone plasmid, lowercase; Insert, uppercase)

| Name | 5'-3' sequence | Note |
|---|---|---|
| Citrine cassette-5F | acgccgttgatgtggacgccgTTACTCTTCTTCTTGATCA | ApaI site of pGWB501 |
| Citrine cassette-3R | gttgaaggagccactcagccGAAAGGGGTTTAGGGTTTAA | SacII site of pGWB501 |
| pGWB-SMXL7pro-5F | ttgcatgcctgcaggtcgactCTATTTTATTTATGTGACAGTTT | XbaI site of pGWB501 |
| pGWB-SMXL7pro-3R | tttttgtacaaacttgttgataactCTAGCGTCGCCGGTTTAGTT | XbaI site of pGWB501 |
| pGWB-SMAX1pro-5F | ttgcatgcctgcaggtcgactAAAAGTAGATTTATTTTGTA | XbaI site of pGWB501 |
| pGWB-SMAX1pro-3R | tttttgtacaaacttgttgataactCGTTCTTCGTTTACTTCCAC | XbaI site of pGWB501 |

II. Primers for cloning Gateway entry clones
(attB, lowercase; overlapped flanking sequences, underlined lowercase; start codons, red; stop codons, blue)

| | | |
|---|---|---|
| BP221-SMAX1-5F | ggggacaagtttgtacaaaaaagcaggctcgATGAGAGCTGGTTTAAGTAC | Amplification of SMAX1 |
| BP221-SMXL7-5F | ggggacaagtttgtacaaaaaagcaggctccATGCCGACACCAGTAACC | Amplification of SMXL7 |
| OE-SMAX1-FLAG-3R | tccttgtaatcgccggatccgccTACTGCCAAAGTAATAGTTGTC | Amplification of SMAX1 with overlapping sequence for FLAG |
| OE-SMXL7-FLAG-3R | tccttgtaatcgccggatccgccGATCACTTCGACTCTCG | Amplification of SMXL7 with overlapping sequence for FLAG |
| GGSG-FLAG-5F | GGCGGATCCGGCGATTACAA | Amplification of FLAG |
| BP221-FLAG-3R | ggggaccactttgtacaagaaagctgggtcCTACTTGTCATCATCATCCTTG TAATC | Amplification of FLAG |
| SMAX1$_{N}$-158-3R | TTTAACGGCGGGACTTGAAAAG | Amplification of SMAX1$_{N}$ |
| OE-SMXLχ71O-5F | gaagcaccgacattaaaGCTACAATTGAACAGTCGTTG | Amplification of SMAX1$_{D1M}$ with overlapping sequence for SMXL7$_{N}$ |
| OE-SMXLχO17-3R | gagagtagttccctgagagaCTTAAACAAATCAATGTCTAACG | Amplification of SMAX1$_{D1M}$ with overlapping sequence for SMXL7$_{D2}$ |
| SMAX1$_{D2}$-611-5F | AAGCTGTTGAAGGGAATGAC | Amplification of SMAX1$_{D2}$ |
| SMXL7$_{N}$-174-3R | TTTAATGTCGGTGCTTCTAAAC | Amplification of SMXL7$_{N}$ |
| OE-SMXLχ17O-5F | cttttcaagtcccgccgttaaaCTCGACGTGCTTCATCCTCC | Amplification of SMXL7$_{D1M}$ with overlapping sequence for SMAX1$_{N}$ |
| OE-SMXLχO71-3R | gtcattcccttcaacagcttCTTGAAATCTTTGCAGTATCTC | Amplification of SMXL7$_{D1M}$ with overlapping sequence for SMAX1$_{D2}$ |
| SMXL7$_{D2}$-630-5F | TCTCTCAGGGAACTACTCTCTC | Amplification of SMXL7$_{D2}$ |
| BP221-SV40 NLS type1-5F | ggggacaagtttgtacaaaaaagcaggctcgATGCCCAAGAAGAAGCGTAA GG | Amplification of SV40 NLS-fused SMAX1$_{N}$ or SMXL7$_{N}$ |
| BP221-SV40NLS type2-5F | ggggacaagtttgtacaaaaaagcaggctcaATGCCAAAAAAGAAGAGAAA GG | Amplification of SV40 NLS-fused SMAX1ΔN or SMXL7ΔN |
| OE-NLS-SMAX1ΔN-5F | aaaggtagaagaccccactagtGCTACAATTGAACAGTCG | Amplification of SV40 NLS-fused SMAX1ΔN |
| OE-NLS-SMXL7ΔN-5F | aaaggtagaagaccccactagtCTCGACGTGCTTCATCCT | Amplification of SV40 NLS-fused SMXL7ΔN |
| BP221-SRDX-stop-3R | ggggaccactttgtacaagaaagctgggtcTCAAGCGAAACCGAGCCTGAG CT | Amplification of SRDX-fused constructs |
| SMXL7ΔRGKT-5F | GACAGTCTTGACGATAGATTCGTTGTTGATTACATTGCTGG | For removal of RGKT motif of SMXL7 |
| SMXL7ΔRGKT-3R | GAATCTATCGTCAAGACTGTC | For removal of RGKT motif of SMXL7 |
| SMAX1ΔRGKT-5F | GAGATGGAAATTCTAGTTTCGCG | For removal of RGKT motif of SMAX1 |
| SMAX1ΔRGKT-3R | GAAACTAGAATTTCCATCTCCAGCATC | For removal of RGKT motif of SMAX1 |
| OE-NLS-GFP-5F | aaaggtagaagaccccactagtATGGTGAGCAAGGGCGAGGA | Amplification of of NLS-GFP constructs |
| OE-GFP-FLAG-3R | tccttgtaatcgccggatccgccCTTGTACAGCTCGTCCATGCC | Amplification of GFP with overlapping sequence for FLAG tag |
| SMAX1$_{N210}$-3R | TTGTACCGACGAAGCGTTCTG | Amplification of SMAX1$_{N210}$ |
| OE-SMXLχ1$_{210}$7O-5F | TCGGTACAAACGTCAAGATCTCGTATTCCTC | Amplification of extended SMXL7$_{D1M}$ with overlapping sequence for SMAX1$_{N210}$ |

| II. Primers for genotyping | | |
|---|---|---|
| smax1-2_LP | GTGGCAACTGTTTAGGCTGAG | Li et al., 2022, paired with LBb1.3[45] |
| smax1-2_RP | AAGCTAGCTTTTCAAGTCCCG | Li et al., 2022, paired with LBb1.3[45] |
| smxl2-1_RP | CCACTTCAGTGTCGAGCTCTC | paired with LB1 |
| smxl2-1_LP | TTGCTCCCAAGCCTAATCAAAAC | paired with LB1 |
| smxl6-4_LP | AGCCAGAGAAAGACTCGAACC | Wang et al., 2015, paired with LBb1.3[40] |
| smxl6-4_RP | TCCGAAATTAAGCTCGATGTG | Wang et al., 2015, paired with LBb1.3[40] |
| smxl7-3_LP | GATCAAGAAACGAACGCTGAG | Wang et al., 2015, paired with WiscDsLox-LB[40] |
| smxl7-3_RP | CGTATTAGCCTCTCGGATTCC | Wang et al., 2015, paired with WiscDsLox-LB[40] |
| smxl8-1_LP | GAATCACAAATTCTGCATGGC | Wang et al., 2015, paired with LBb1.3[40] |
| smxl8-1_RP | CTGACGAAGCTCCACTTTCAC | Wang et al., 2015, paired with LBb1.3[40] |
| max3-9-F | GGTCACTTGCAACGCTGAAG | *max3-9* 102bp; MAX3 75bp |
| max3-9 -R | GAATTAAAAGATTATTTACCACAAAATGTGAAGTTGCT | *max3-9* 102bp; MAX3 75bp |
| SAIL_LB1 | GCCTTTTCAGAAATGGATAAATAGCCTTGCTTCC | |
| SALK_LBb1.3 | ATTTTGCCGATTTCGGAAC | |
| WiscDsLox-LB-p745 | AACGTCCGCAATGTGTTATTAAGTTGTC | |

### References


1. Pruneda-Paz, J. L. *et al.* A genome-scale resource for the functional characterization of Arabidopsis transcription factors. *Cell Rep.* **8**, 622–632 (2014).

2. Verma, S., Attuluri, V. P. S. & Robert, H. S. Transcriptional control of Arabidopsis seed development. *Planta* **255**, 90 (2022).

3. Ali, F., Qanmber, G., Li, F. & Wang, Z. Updated role of ABA in seed maturation, dormancy, and germination. *J. Advert. Res.* **35**, 199–214 (2022).

4. Shibata, M. & Sugimoto, K. A gene regulatory network for root hair development. *J. Plant Res.* **132**, 301–309 (2019).

5. Humphreys, J. L., Beveridge, C. & Tanurdzic, M. Strigolactone-dependent gene regulation requires chromatin remodeling. *bioRxiv* 2023.02.25.529999 (2023) doi:10.1101/2023.02.25.529999.

6. Shi, H. *et al.* Genome-wide regulation of light-controlled seedling morphogenesis by three families of transcription factors. *Proc. Natl. Acad. Sci. U. S. A.* **115**, 6482–6487 (2018).

7. Wang, L. *et al.* Transcriptional regulation of strigolactone signalling in Arabidopsis. *Nature* **583**, 277–281 (2020).

8. Pucciariello, O. *et al.* Rewiring of auxin signaling under persistent shade. *Proc. Natl. Acad. Sci. U. S. A.* **115**, 5612–5617 (2018).

9. Hu, J., Ji, Y., Hu, X., Sun, S. & Wang, X. BES1 Functions as the Co-regulator of D53-like SMXLs to Inhibit BRC1 Expression in Strigolactone-Regulated Shoot Branching in Arabidopsis. *Plant Commun* **1**, 100014 (2020).

10. Cao, J. *et al.* Multi-layered roles of BBX proteins in plant growth and development. *Stress Biol* **3**, 1 (2023).

11. Hou, Q. *et al.* Overexpression of HLH4 Inhibits Cell Elongation and Anthocyanin Biosynthesis in Arabidopsis thaliana. *Cells* **11**, (2022).

12. Jiao, Y., Lau, O. S. & Deng, X. W. Light-regulated transcriptional networks in higher plants. *Nat. Rev. Genet.* **8**, 217–230 (2007).

13. Ju, L. *et al.* JAZ proteins modulate seed germination through interaction with ABI5 in bread wheat and Arabidopsis. *New Phytol.* **223**, 246–260 (2019).

14. Wang, P. *et al.* Photomorphogenesis in plants: The central role of phytochrome interacting factors (PIFs). *Environ. Exp. Bot.* **194**, 104704 (2022).

15. Jia, K.-P., Luo, Q., He, S.-B., Lu, X.-D. & Yang, H.-Q. Strigolactone-regulated hypocotyl elongation is dependent on cryptochrome and phytochrome signaling pathways in Arabidopsis. *Mol. Plant* **7**, 528–540 (2014).

16. Park, Y.-J., Kim, J. Y. & Park, C.-M. SMAX1 potentiates phytochrome B-mediated hypocotyl thermomorphogenesis. *Plant Cell* **34**, 2671–2687 (2022).

17. Burko, Y. *et al.* PIF7 is a master regulator of thermomorphogenesis in shade. *Nat. Commun.* **13**, 4942 (2022).

18. Ding, J., Zhang, B., Li, Y., André, D. & Nilsson, O. Phytochrome B and PHYTOCHROME INTERACTING FACTOR8 modulate seasonal growth in trees. *New Phytol.* **232**, 2339–2352 (2021).

19. Bian, Y. *et al.* PIFs- and COP1-HY5-mediated temperature signaling in higher plants. *Stress Biol* **2**, 35 (2022).

20. Yadav, A., Ravindran, N., Singh, D., Rahul, P. V. & Datta, S. Role of Arabidopsis BBX proteins in light signaling. *J. Plant Biochem. Biotechnol.* **29**, 623–635 (2020).

21. Ali, S., Khan, N. & Xie, L. Molecular and Hormonal Regulation of Leaf Morphogenesis in Arabidopsis. *Int. J. Mol. Sci.* **21**, (2020).

22. Romanova, M. A., Domashkina, V. V., Maksimova, A. I., Pawlowski, K. & Voitsekhovskaja, O. V. All together now: Cellular and molecular aspects of leaf development in lycophytes, ferns, and seed plants. *Frontiers in Ecology and Evolution* **11**, (2023).

23. Jin, H. & Zhu, Z. HOOKLESS1 is a positive regulator in Arabidopsis thermomorphogenesis. *Sci. China Life Sci.* **62**, 423–425 (2019).

24. Zhao, H., Zhang, Y. & Zheng, Y. Integration of ABA, GA, and light signaling in seed germination through the regulation of ABI5. *Front. Plant Sci.* **13**, 1000803 (2022).

25. Sun, H. *et al.* Strigolactone and gibberellin signaling coordinately regulate metabolic adaptations to changes in nitrogen availability in rice. *Mol. Plant* **16**, 588–598 (2023).

26. Capella, M., Ribone, P. A., Arce, A. L. & Chan, R. L. Arabidopsis thaliana HomeoBox 1 (AtHB1), a Homedomain-Leucine Zipper I (HD-Zip I) transcription factor, is regulated by PHYTOCHROME-INTERACTING FACTOR 1 to promote hypocotyl elongation. *New Phytol.* **207**, 669–682 (2015).

27. Jiang, Z., Xu, G., Jing, Y., Tang, W. & Lin, R. Phytochrome B and REVEILLE1/2-mediated signalling controls seed dormancy and germination in Arabidopsis. *Nat. Commun.* **7**, 12377 (2016).

28. Song, X. *et al.* IPA1 functions as a downstream transcription factor repressed by D53 in strigolactone signaling in rice. *Cell Res.* **27**, 1128–1141 (2017).

29. Hwang, Y. *et al.* Anterograde signaling controls plastid transcription via sigma factors separately from nuclear photosynthesis genes. *Nat. Commun.* **13**, 7440 (2022).

30. Challa, K. R., Aggarwal, P. & Nath, U. Activation of YUCCA5 by the Transcription Factor TCP4 Integrates Developmental and Environmental Signals to Promote Hypocotyl Elongation in Arabidopsis. *Plant Cell* **28**, 2117–2130 (2016).

31. Challa, K. R., Rath, M. & Nath, U. The CIN-TCP transcription factors promote commitment to differentiation in Arabidopsis leaf pavement cells via both auxin-dependent and independent pathways. *PLoS Genet.* **15**, e1007988 (2019).

32. Muhr, M., Paulat, M., Awwanah, M., Brinkkötter, M. & Teichmann, T. CRISPR/Cas9-mediated knockout of Populus BRANCHED1 and BRANCHED2 orthologs reveals a major function in bud outgrowth control. *Tree Physiol.* **38**, 1588–1597 (2018).

33. Rinne, P. L. H. *et al.* Long and short photoperiod buds in hybrid aspen share structural development and expression patterns of marker genes. *J. Exp. Bot.* **66**, 6745–6760 (2015).

34. Hur, Y.-S. *et al.* Arabidopsis transcription factor TCP13 promotes shade avoidance syndrome-like responses by directly targeting a subset of shade-responsive gene promoters. *J. Exp. Bot.* **75**, 241–257 (2024).

35. Zhou, Y. *et al.* TCP Transcription Factors Associate with PHYTOCHROME INTERACTING FACTOR 4 and CRYPTOCHROME 1 to Regulate Thermomorphogenesis in Arabidopsis thaliana. *iScience* **15**, 600–610 (2019).

36. Ferrero, L. V., Gastaldi, V., Ariel, F. D., Viola, I. L. & Gonzalez, D. H. Class I TCP proteins TCP14 and TCP15 are required for elongation and gene expression responses to auxin. *Plant Mol. Biol.* **105**, 147–159 (2021).

37. Zhang, W. *et al.* The MPK8-TCP14 pathway promotes seed germination in Arabidopsis. *Plant J.* **100**, 677–692 (2019).

38. Viola, I. L., Reinheimer, R., Ripoll, R., Manassero, N. G. U. & Gonzalez, D. H. Determinants of the DNA binding specificity of class I and class II TCP transcription factors. *J. Biol. Chem.* **287**, 347–356 (2012).

39. Aguilar-Martínez, J. A., Poza-Carrión, C. & Cubas, P. Arabidopsis BRANCHED1 acts as an integrator of branching signals within axillary buds. *Plant Cell* **19**, 458–472 (2007).

40. Wang, L. *et al.* Strigolactone Signaling in Arabidopsis Regulates Shoot Development by Targeting D53-Like SMXL Repressor Proteins for Ubiquitination and Degradation. *Plant Cell* **27**, 3128–3142 (2015).

41. Braun, N. *et al.* The pea TCP transcription factor PsBRC1 acts downstream of Strigolactones to control shoot branching. *Plant Physiol.* **158**, 225–238 (2012).

42. Danisman, S. *et al.* Analysis of functional redundancies within the Arabidopsis TCP transcription factor family. *J. Exp. Bot.* **64**, 5673–5685 (2013).

43. Han, X. *et al.* Arabidopsis Transcription Factor TCP5 Controls Plant Thermomorphogenesis by Positively Regulating PIF4 Activity. *iScience* **15**, 611–622 (2019).

44. Shen, N., Hou, S., Tu, G., Lan, W. & Jing, Y. Transcription Factor WRKY33 Mediates the Phosphate Deficiency-Induced Remodeling of Root Architecture by Modulating Iron Homeostasis in Arabidopsis Roots. *Int. J. Mol. Sci.* **22**, (2021).

45. Li, Q. et al. The strigolactone receptor D14 targets SMAX1 for degradation in response to GR24 treatment and osmotic stress. Plant Commun 3, 100303 (2022).
